# Single-cell transcriptional mapping reveals genetic and non-genetic determinants of aberrant differentiation in AML

**DOI:** 10.1101/2023.12.26.573390

**Authors:** Andy G.X. Zeng, Ilaria Iacobucci, Sayyam Shah, Amanda Mitchell, Gordon Wong, Suraj Bansal, David Chen, Qingsong Gao, Hyerin Kim, James A. Kennedy, Andrea Arruda, Mark D. Minden, Torsten Haferlach, Charles G. Mullighan, John E. Dick

## Abstract

In acute myeloid leukemia (AML), genetic mutations distort hematopoietic differentiation, resulting in the accumulation of leukemic blasts. Yet, it remains unclear how these mutations intersect with cellular origins and whether they converge upon similar differentiation patterns. Single-cell RNA sequencing (scRNA-seq) has enabled high-resolution mapping of the relationship between leukemia and normal cell states, yet this application is hampered by imprecise reference maps of normal hematopoiesis and small sample sizes among patient cohorts. As a first step we constructed a reference atlas of human bone marrow hematopoiesis from 263,519 single-cell transcriptomes spanning 55 cellular states, that was benchmarked against independent datasets of immunophenotypically pure hematopoietic stem and progenitor cells. Using this reference atlas, we mapped over 1.2 million single-cell transcriptomes spanning 318 AML, mixed phenotype acute leukemia (MPAL), and acute erythroid leukemia (AEL) samples. This large-scale analysis, together with systematic mapping of genotype-to-phenotype associations between driver mutations and differentiation landscapes, revealed convergence of diverse genetic alterations on twelve recurrent patterns of aberrant differentiation in AML. This included unconventional lymphoid and erythroid priming linked to *RUNX1* and *TP53* mutations, respectively. We also identified non-genetic determinants of AML differentiation such as two subgroups of *KMT2A*-rearranged AML that differ in the identity of their leukemic stem cells (LSCs), likely reflecting distinct cellular origins. Furthermore, distinct LSC-driven hierarchies can co-exist within individual patients, providing insights into AML evolution. Together, precise mapping of normal and malignant cell states provides a framework for advancing the study and disease classification of hematologic malignancies thereby informing therapy development.

## Introduction

Hematopoietic stem and progenitor cells (HSPCs) traverse complex differentiation landscapes, following well-defined trajectories in order to produce mature blood cells^1^. In acute myeloid leukemia (AML), genetic mutations distort these differentiation trajectories through inappropriate induction of aberrant self-renewal, lineage biases, or differentiation blocks. While many studies have evaluated how specific AML driver mutations perturb hematopoiesis^2–6^, a comprehensive understanding of the genotype-to-phenotype relationships across a wide spectrum of AML cases remains elusive. This also raises an important question of whether the consequences of these perturbations to hematopoietic differentiation converge on recurrent phenotypes or whether they are lesion-specific. Addressing this could provide deeper insight into the pathogenesis of each patient’s disease and may improve the design of differentiation therapies.

In clinical practice, leukemic blasts are assigned to broad hematopoietic lineages based on morphologic and immunophenotypic features of normal hematopoiesis^7^. Yet, these features lack the precision in discerning between the specific cell states that reside within a hematopoietic lineage, particularly at the level of immature blasts. In contrast, scRNA-seq enables precise determination of leukemia cell state using thousands of gene expression features. This is particularly relevant for acute myeloid leukemia (AML) wherein leukemic cells form cellular hierarchies spanning multiple differentiation stages, sustained by leukemia stem cells (LSCs) at the apex. Accordingly, scRNA-seq has now been applied extensively to map cellular hierarchies in normal hematopoiesis^8–13^ as well as AML^14–28^, revealing variation in AML hierarchy composition that has been linked to survival, relapse, and drug sensitivity^29,30^. Yet, these studies have often been constrained by the use of early reference maps that provide a limited view of normal hematopoietic differentiation, owing to low cell numbers or insufficient representation of HSPCs. Along with small sample sizes of AML patient samples, these limitations have hindered a comprehensive understanding of the diverse patterns in which differentiation is disrupted across a spectrum of AML cases.

In this study, we constructed a single-cell reference atlas of human hematopoietic differentiation and mapped single-cell transcriptomes from more than 300 myeloid leukemia patient samples. This revealed phenotypic convergence upon 12 recurrent patterns of aberrant differentiation, linked to genetic driver alterations as well as differences in cellular origins and LSC identities. Together, this establishes a new framework for understanding aberrant differentiation in AML.

## Methods

### Biological Samples

All samples were collected with informed consent according to procedures approved by either the University Health Network (UHN) Research Ethics Board or St. Jude Children’s Research Hospital (SJCRH) Institutional Review Board. Human CB samples were obtained from Trillium Health, Credit Valley, and William Osler Hospitals in Ontario. Primary AML samples obtained from the Munich Leukemia Laboratory (MLL) (n = 12) were processed for scRNA-seq with 10x 5’ v1.1 technology at SJCRH, and 2 primary AML samples obtained from the Princess Margaret Hospital (PMH) Leukaemia Bank were processed for scRNA-seq with 10x 3’ v3 technology at PMH, per manufacturer instructions. Clinical data for leukemia samples in this study are outlined in **supplemental Tables 5 and 14**. Normal datasets used to develop the Bone Marrow Reference Map, along with additional normal and leukemic samples analyzed by scRNA-seq data from published studies, are outlined in **supplemental Table 4**.

### Computational Analyses

scRNA-seq preprocessing was performed using seurat^31^ and scanpy^32^ and each dataset was filtered based on their quality control (QC) metric distributions. Batch correction of scRNA-seq datasets was performed with harmony^33^ and reference map projection was performed using symphony^34^. Differential expression analyses utilized DESeq2^35^ at the level of pseudo-bulk profiles. Gene expression modules were defined by cNMF^36^, transcription factor regulons were defined by pySCENIC^37^, and signature scoring was performed using AUCell^38^. Pseudo-time inference was performed using Monocle3^39^. For genotyping analysis from scRNA-seq, expressed mutations and gene fusions were called by cb_sniffer (https://github.com/genome/cb_sniffer) while copy number alterations were called by inferCNV (https://github.com/broadinstitute/inferCNV).

Full details regarding additional experimental and computational methods used in this study are provided in the supplemental Methods.

## Results

### A single-cell reference landscape of human bone marrow hematopoiesis

We sought to develop a high quality scRNA-seq reference atlas of bone marrow hematopoiesis to facilitate analysis of scRNA-seq data from benign and malignant hematopoiesis. To ensure balanced proportions of rare HSPCs alongside terminally differentiated populations, we integrated two scRNA-seq datasets from unsorted bulk BM cells^40–42^ with three scRNA-seq datasets of CD34+ BM cells^43–45^ and one scRNA-seq dataset comprising data from both bulk and CD34+ BM cells^46^. After quality control, our analysis included 263,159 high-quality single-cell transcriptomes from BM cells wherein 89,404 cells were derived from CD34+ sorted BM samples and 173,755 cells were derived from unsorted bulk BM samples. Following dimensionality reduction and focused clustering (supplemental Methods), we identified 55 hematopoietic cell states (**Figure 1A, supplemental Tables 1 and 2**). Within these, 40 states spanned continuous differentiation trajectories that begin from primitive HSCs and end at terminal states spanning erythroblasts (Ery), megakaryocytes (Mk), granulocytes (Gran), monocytes (Mono), conventional dendritic cells (cDC), plasmacytoid dendritic cells (pDC), and B cells. The remaining 15 cell states were positioned outside of this differentiation hierarchy, encompassing T, NK, plasma cell, and stromal cell populations.

**Figure 1.**
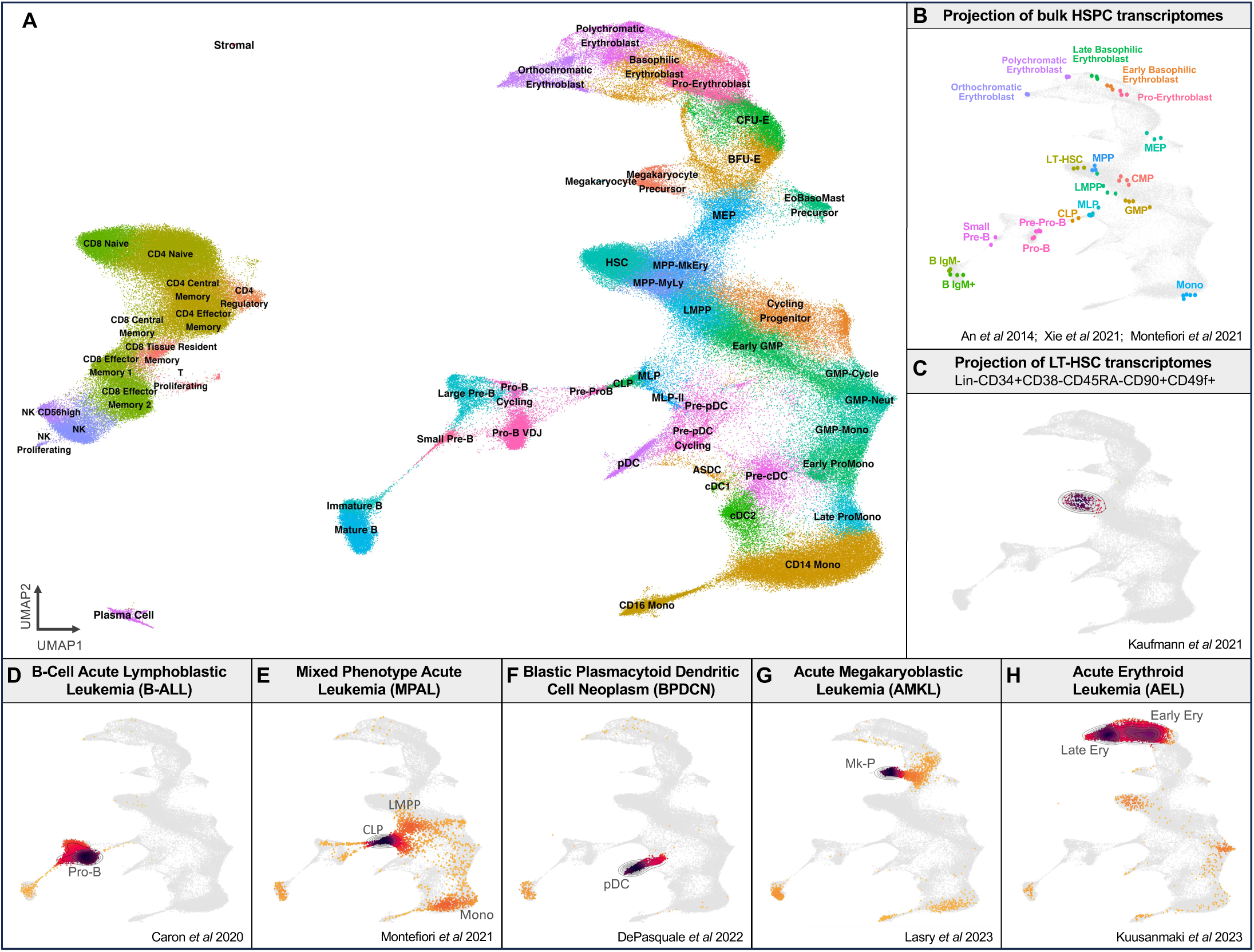
A single-cell reference landscape of human bone marrow hematopoiesis. **A)** Single-cell transcriptional atlas of human hematopoiesis with balanced representation of CD34+ stem and progenitor cells together with terminally differentiated populations, comprising 263,159 bone marrow cells spanning 55 cell states across 45 donors from six studies. **B)** Projection of bulk transcriptomes from purified hematopoietic cell populations onto BoneMarrowMap. **C)** Projection of single-cell transcriptomes from immunophenotypic LT-HSCs (Lin-CD34+CD38-CD45RA-CD90+CD49f+). **D-H)** Projection of scRNA-seq profiles from leukemia samples spanning multiple diagnoses, including **D)** B cell acute lymphoblastic leukemia (B-ALL), **E)** mixed phenotype acute leukemia (MPAL), **F)** blastic plasmacytoid dendritic cell neoplasm (BPDCN), **G)** acute megakaryoblastic leukemia (AMKL), and **H)** acute erythroid leukemia (AEL).

To validate the reference map and cell state assignments, transcriptional cell state annotations were compared against functionally-defined HSPC populations. Transcriptional projection of bulk RNA-seq profiles from flow-sorted populations along the hematopoietic hierarchy^47–49^ was consistent with their ordering along our UMAP embedding of hematopoiesis (**Figure 1B**). Furthermore, projection of scRNA-seq profiles from purified long-term (LT) HSC^8,9,12,50–52^ as well as purified hematopoietic progenitor populations^12,53,54^ including multi-potent progenitors (MPPs), megakaryocyte-erythrocyte progenitors (MEPs), lympho-myeloid primed progenitors (LMPPs), multi-lymphoid progenitors (MLPs), and granulocyte-monocyte progenitors (GMPs), showed broad agreement between transcriptionally-defined cell states and functionally-defined HSPC fractions, including high concordance between transcriptional and immunophenotypic HSCs (**Figure 1C, supplemental Note 1, supplemental Figures 1-2**).

### Uncovering gene expression programs across hematopoietic differentiation

To leverage this transcriptome-level expression information across 55 cell states, we sought to identify gene expression programs underlying hematopoietic differentiation in an unsupervised manner. Consensus NMF^36^ was performed across the reference atlas, resulting in the identification of 48 distinct gene expression programs, each representing groups of genes with correlated expression across human hematopoiesis: 36 programs were specific to hematopoietic cell states, whereas 12 programs captured the activity of biological processes spanning the cell cycle, oxidative phosphorylation, and stress response, among others (**supplemental Figure 3A, supplemental Table 3**). To characterize these programs, we identified key transcription factor (TF) regulons (**supplemental Figure 3B**) and biological pathways (**supplemental Figure 3C**) underlying these molecular programs. Notably, while the G2 and M phases of the cell cycle are typically grouped into a single molecular program^55^, NMF decoupled these stages into distinct programs revealing stepwise activation of S-phase, G2-phase, and M-phase programs within our dataset (**supplemental Figure 3D-K**).

To understand how these gene expression programs vary across the continuum of human hematopoietic differentiation, we performed pseudo-time analysis^39^ along each lineage captured in our reference map. We also developed an approach to transfer pseudo-time estimates onto query data, which we validated in an independent dataset^10^ (**supplemental Figure 4A-C**). Pseudo-time analyses revealed key genes and transcription factors underlying successive waves of molecular programs induced during hematopoietic differentiation along human monocyte and erythroid lineages (**supplemental Figure 4D-I**) as well as B cell, pDC, cDC (**supplemental Figure 5A-I**), megakaryocyte, eosinophil/basophil/mast cell (EoBasoMast), and neutrophil lineages (**supplemental Figure 6A-I**). Collectively, this provides a valuable resource for exploring the molecular programs which underlie the early and late lineage transitions in human hematopoietic differentiation.

### Assignment of individual leukemia cells to precise hematopoietic states

Having developed a comprehensive reference atlas of human hematopoiesis, we asked whether this resource could be applied to better understand how hematopoietic differentiation becomes corrupted in acute leukemia. Phenotypic classification of acute leukemia involves the determination of hematopoietic lineage through a small number of cell surface markers, thus we reasoned that scRNA-seq projection using thousands of gene expression features onto our hematopoietic cell atlas may improve lineage classification in acute leukemia. To evaluate our transcriptional approach for leukemia cell classification, we projected scRNA-seq profiles from leukemia patient samples with clearly defined lineage features. As expected, Pro-B cells were enriched in B-acute lymphoblastic leukemia (B-ALL)^56^ (**Figure 1D**), multipotent MLPs were enriched in mixed-phenotype acute leukemia (MPAL)^48^ (**Figure 1E**), pDCs were enriched in blastic plasmacytoid dendritic cell neoplasm (BPDCN)^57^ (**Figure 1F**), Mk precursors (MkP) were enriched in acute megakaryoblastic leukemia (AMKL)^20^ (**Figure 1G**), and erythroblasts were enriched in acute erythroid leukemia (AEL)^58^ (**Figure 1H**).

To map AML samples to precise cellular states across human hematopoietic differentiation, we first performed scRNA-seq profiling on 12 AML patient samples harboring either myelodysplasia (MDS)-related changes or classical genetic alterations in *NPM1* or *KMT2A*. Next, we re-analyzed single-cell transcriptomes from twenty additional studies^14,15,18–26,46,48,58–64^ (151 adult AML samples, 136 pediatric AML samples, and 18 MPAL samples), amounting to a total of 318 patient samples (**supplemental Tables 4 and 5**). From these patient samples, 1,223,411 cells were confidently mapped to hematopoietic cell states. After exclusion of non-leukemic mature lymphocytes, we performed composition analysis of leukemia cells mapped to 38 cell states (**Figure 2A, supplemental Figure 7A, supplemental Table 6**). Specifically, similar cell states that co-occurred across patient samples were collapsed together, thus consolidating 38 precise cell states into 13 broad differentiation stages spanning HSC/MPP, LMPP, Early Lymphoid, MEP/MkP, EoBasoMast, Early Erythroid, Late Erythroid, GMP, ProMono, Mono, and cDC (**supplemental Figure 7B, supplemental Tables 7 and 8**).

**Figure 2.**
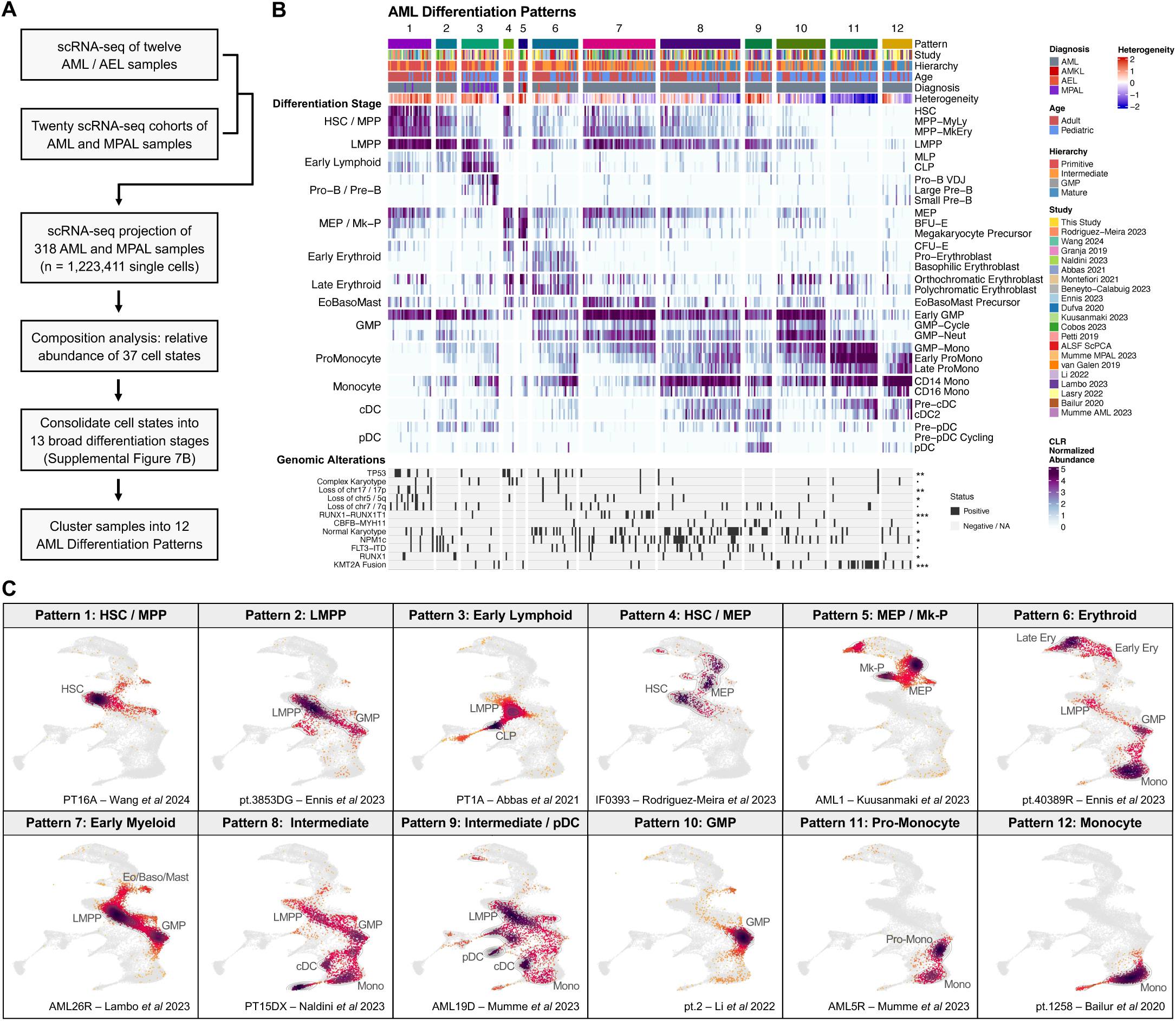
Projection and classification of single cell transcriptomes in acute leukemia. **A)** Cell state projection and composition analysis of scRNA-seq profiles from 318 acute myeloid leukemia (AML) and MPAL samples to identify recurrent patterns of AML differentiation. **B)** Twelve AML differentiation patterns identified through cell state projection and composition analysis. The heatmap depicts the centered log ratio normalized abundance of individual cell states within each patient sample, clipped at 0 and 4.5 for visualization purposes. These cell states are grouped into broader differentiation stages displayed on the left of the heatmap. Each sample is annotated based on the study of origin, AML hierarchy classification from deconvolution of pseudo-bulk RNA profiles, pediatric or adult age group, and specific disease diagnosis spanning AML, MPAL, AMKL, and AEL. A measure of cell state heterogeneity is also depicted for each patient sample, quantified by the Shannon diversity index. Key genomic alterations are also annotated for each scRNA-seq patient sample with significance values derived through chi-squared tests (• P<0.10; * P<0.05; ** P<0.01; *** P<0.001). **C)** Projection results of representative patient samples for each of the twelve AML differentiation patterns.

Projected AML cells retained gene expression programs which reflected their assigned stages along hematopoietic differentiation trajectories (**supplemental Figure 7C-D**). Importantly, the expression of common therapeutic targets in AML varied dramatically by differentiation stage, highlighting the relevance of cellular heterogeneity for understanding drug response in AML (**supplemental Figure 8A**). Further, LSC signatures (DE genes from sorted LSC+ fractions^65^ as well as a Quiescent LSPC program^29^) were enriched across multiple stem and progenitor differentiation stages, with this being most pronounced at the HSC/MPP and LMPP stages (**supplemental Figure 8B-C**). Accordingly, Quiescent LSPC, a state previously linked to functional LSC potential^29^, localized primarily to HSC/MPP and LMPP and sometimes Early Lymphoid and MEP in a patient-specific manner^14,18,26,29^ (**supplemental Figure 8D**), suggesting that LSC potential may reside within distinct stem and progenitor compartments in different patients.

Next, we sought to compare the differentiation landscapes of the 318 patient samples. Analysis of leukemia cell composition revealed twelve recurrent patterns of AML differentiation (**Figure 2B-C, supplemental Figure 9, supplemental Table 5**). Patterns 1 and 2 were defined by early blocks in differentiation, resulting in accumulation of HSC/MPP (p1) and LMPP (p2) populations. Patterns 3 - 7 featured enrichment of hematopoietic progenitors from distinct lineages, such as Early Lymphoid (p3), MEP/Mk-P (p4, p5), Erythroid (p6), and Early Myeloid (p7). Patterns 8 and 9 featured disease involvement along each stage of monocytic, cDC, and pDC (p9) differentiation, lacking obvious differentiation blocks along those trajectories. Patterns 10-12 were specifically enriched for committed myeloid progenitors including GMP (p10), ProMono (p11), and Mono (p11, p12).

Notably, differentiation patterns that were most enriched for primitive HSC/MPP (p1 and p4) were under-represented among pediatric AML compared to adult AML (2% vs 20% at diagnosis, *P*=6e-6). We confirmed that the proportion of leukemic cells mapping HSC and MPP states was lower in pediatric AML (*P*=9e-9). Alternatively, differentiation patterns 11 (ProMono) and 12 (Mono) were over-represented in pediatric AML compared to adult AML (28% vs 12% at diagnosis, *P*=0.001). In these latter subgroups many patient samples lack detectable HSC/MPP or LMPP, likely reflecting differences in their cellular origins. Thus, despite the diverse genetic mechanisms by which hematopoietic differentiation can be disrupted, these data suggest considerable convergence upon twelve recurrent patterns of aberrant differentiation in AML.

### Genetic determinants of lineage biases in acute myeloid leukemia

Next, we asked whether these differentiation patterns in AML were associated with the genetic driver alterations that underlie each patient’s disease. We found that a specific differentiation pattern could arise from multiple genetic drivers (**Figure 2B**). We next sought to utilize bulk transcriptomes at the scale of patient cohorts to perform a more comprehensive characterization of genotype-to-phenotype associations in AML. To this end, differentially expressed marker genes were identified for each AML differentiation stage (**supplemental Figure 10A-B**) and used to train sparse regression models to quantify the abundance of each AML differentiation stage from bulk RNA-seq profiles (supplemental Methods; **supplemental Figure 10C-D, supplemental Table 9**). Leveraging bulk transcriptomes of 1,224 AML patients spanning five patient cohorts^30,66–70^ (**supplemental Table 10**), we mapped the associations between genetic driver alteration status and inferred abundance of each AML differentiation stage (**Figure 3A-B**). In addition to capturing phenotypic convergence between genetic drivers, this allowed us to identify characteristic differentiation landscapes for common driver alterations in AML (**Figure 3C, supplemental Table 11**). These analyses also revealed the specific enrichment patterns for each differentiation stage across existing classification schemes for AML heterogeneity, including blast morphology^71^, WHO 2022^7^, and ICC 2022^72^, AML hierarchy classes^29^, transcriptome-based clusters^70^, and disease subtypes defined through integration of hierarchy, transcriptome, and methylation profiles^73^, among others (**supplemental Figure 11**).

**Figure 3.**
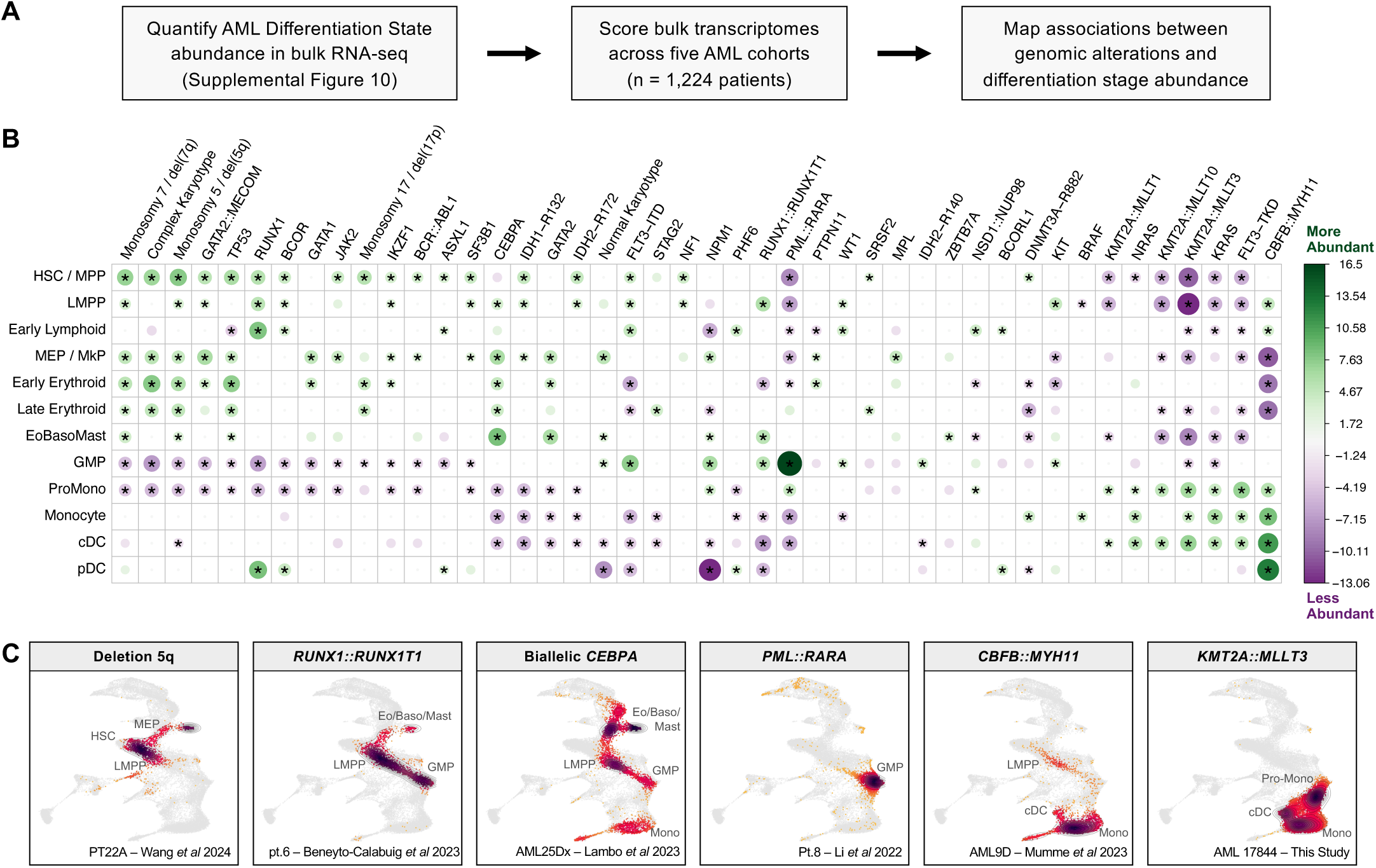
Associations between genetic driver alterations and AML differentiation. **A)** Mapping the relationship of genomic alterations with AML differentiation through bulk transcriptomic quantification of AML differentiation stages across 1,224 AML patient samples from five cohorts re-analyzed by Severens *et al* 2024 (TCGA, BeatAML2, Leucegene, TARGET, LUMC). **B)** Association between inferred abundance of AML differentiation stages with genomic alterations across 1,224 adult and pediatric AML samples profiled by RNA-seq. The strength of each association, quantified as the test statistic from a generalized linear model adjusting for cohort as a covariate, is depicted through the size and color intensity of each dot, with green denoting higher abundance and purple denoting lower abundance. Only associations with unadjusted *P* < 0.05 are shown, associations with FDR < 0.05 are starred. **C)** Projection results of representative patient samples for common genomic alterations in AML.

We next sought to investigate the determinants of unconventional lineage priming in AML. Based on single-cell composition analysis, Pattern 3 consisted of 25 samples enriched for LMPP and Early Lymphoid states and featured co-clustering of 14 MPAL and 11 AML samples (**Figure 4A**), challenging traditional diagnostic boundaries between these two disease classes^7^. This included an AML patient that exhibited an immunophenotypic lineage switch from myeloid to lymphoid from diagnosis to relapse^25^. In this patient, scRNA-seq projection revealed pre-existing lymphoid states at diagnosis that were not detected by immunophenotype, explaining the apparent lineage switch observed at relapse (**Figure 4B**). At the bulk level, *RUNX1* mutations (n=122), which are commonly observed in MPAL^74^, exhibited the strongest association with Early Lymphoid abundance (FDR=1e-13). This association was even more pronounced for biallelic *RUNX1* mutations (**Figure 4C-D**) compared to monoallelic *RUNX1* mutations, suggestive of a dose-dependent relationship. Notably, pDCs have been shown to descend from lymphoid progenitors^75^ and *RUNX1* mutations were also strongly associated with pDC abundance (FDR=5e-15) (**Figure 4C-D**), a finding that is also in concordance with the high frequency of *RUNX1* mutations reported in pDC-AML^76,77^.

**Figure 4.**
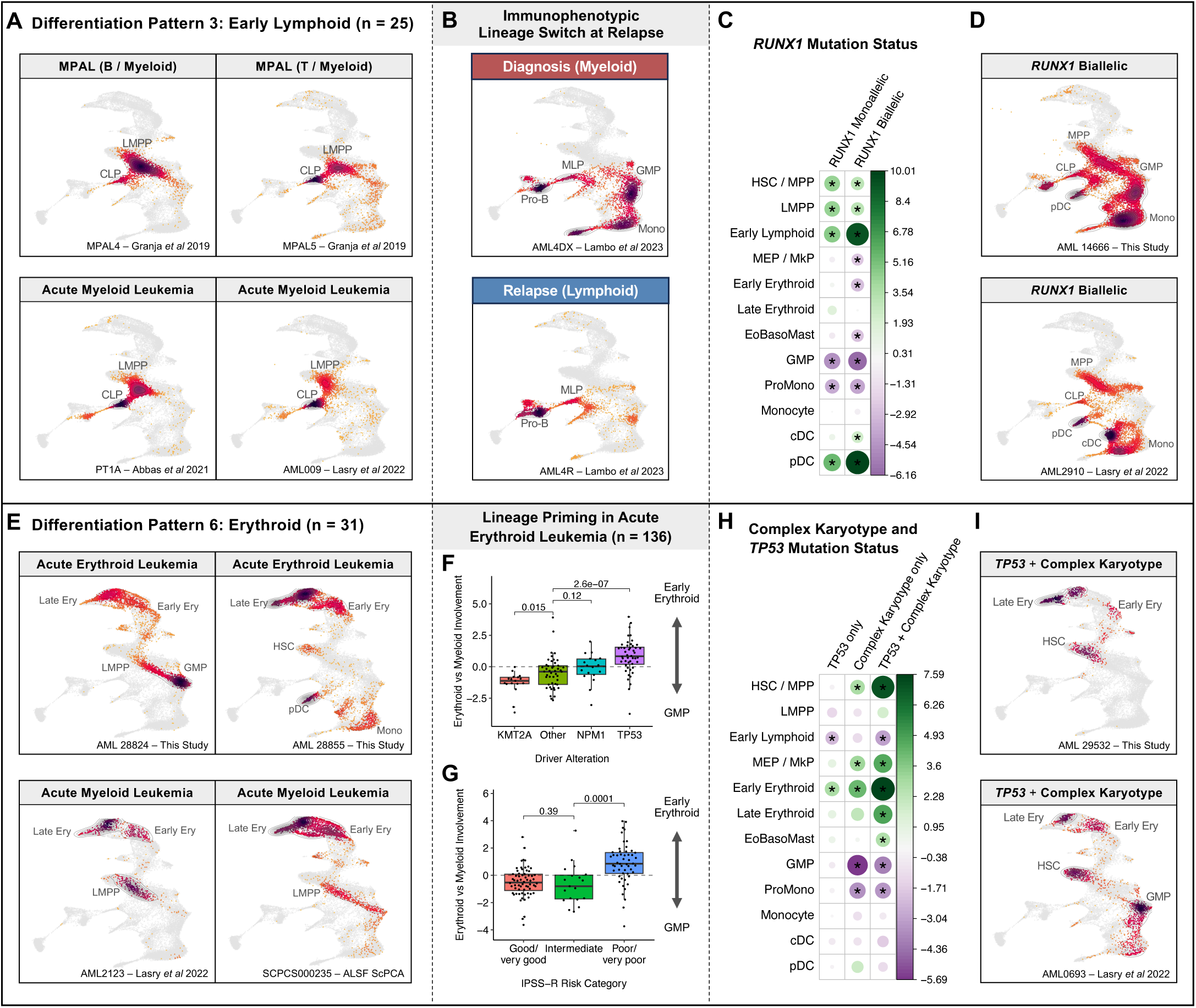
Genetic determinants of lymphoid and erythroid priming in AML. **A)** Representative AML and MPAL patient samples from Differentiation Pattern 3, enriched for Early Lymphoid states. **B)** Example of an infant AML patient sample carrying a *KMT2A::AFF3* fusion, belonging to Differentiation Pattern 3, that experienced an immunophenotypic lineage switch from myeloid to lymphoid from initial diagnosis to disease relapse. **C)** Association between inferred abundance of AML differentiation stages with *RUNX1* mutations (n=87 monoallelic; n=32 biallelic) across 1,224 AML patients profiled by RNA-seq. The strength of each association, quantified as the test statistic from a generalized linear model adjusting for cohort as a covariate, is depicted through the size and color intensity of each dot, wherein green denotes higher abundance and purple denotes lower abundance. Associations with FDR < 0.05 are starred. **D)** Representative patient samples for AML with biallelic *RUNX1* mutations. **E)** Representative AML and AEL patient samples from Differentiation Pattern 6, enriched for Early Erythroid states. **F-G)** Erythroid vs myeloid involvement within 136 AEL samples, depicted as the difference between the inferred Early Erythroid abundance and GMP abundance in each sample. These are depicted in association with **F)** driver mutation status and **G)** clinical risk category. **H)** As with (C), instead depicting the associations of AML differentiation stages with *TP53* only (n=16), Complex Karyotype only (n=95), or *TP53* + Complex Karyotype (n=85) mutation status compared to wildtype samples. **D)** Representative patient samples for AML with Complex Karyotype and *TP53* mutations.

Pattern 6 consisted of 28 AML samples co-clustering with 3 AEL samples based on enrichment for Early Erythroid (CFU-E, Pro-Erythroblast, and Basophilic Erythroblast) states (**Figure 4E**). Across bulk AML transcriptomes, higher Early Erythroid abundance was most enriched among samples with *TP53* mutations (n=101; FDR=8e-12), complex karyotype (n=191; FDR=4e-13), and MDS-related cytogenetic alterations including monosomy 5/del(5q) (n=86; FDR=4e-7), monosomy 7/del(7q) (n=97; FDR=5e-6), and monosomy 17/del(17p) (n=62; FDR=0.0001). *TP53* is the most commonly mutated gene in AEL^78^, and even within AEL samples (n=136, **supplemental Table 12**) we observed greater erythroid involvement in patients with *TP53* mutations (*P*=3e-7) as well as with clinically poor risk disease (*P*=0.0001) (**Figure 4G**). Alterations in *TP53* frequently co-occur with the cytogenetic alterations that also promote erythroid fate, and indeed among AML samples we find that the combination of *TP53* mutations with complex karyotype is more strongly associated with Early Erythroid abundance than either alteration alone (**Figure 4H-I)**.

Beyond the unconventional patterns of lymphoid and erythroid lineage priming, these examples involving *RUNX1* and *TP53* carry principles that can be applied to other genetic alterations. For example, a dose-dependent relationship was also observed for biallelic *CEBPA* mutations, where progenitor involvement was more pronounced at the expense of HSC/MPPs, especially when the favorable-risk bZIP domain was involved^79^ (**supplementary Figure 11H**). Additionally, specific combinations of genetic mutations, particularly those involving *FLT3* and *NPM1,* were capable of producing highly divergent differentiation landscapes (**supplementary Figure 11I-J**). Collectively, this analysis highlights the nuances of genotype-to-phenotype relationships in AML.

### Genetic subclones can impact AML cell hierarchy composition

Recognizing that genomic drivers are associated with lineage priming in AML patients, we next asked whether these relationships could be identified at the level of genetic subclones within individual patients. To answer this question, we employed a variety of analytical approaches, including the identification of expressed mutations or copy number alterations within a subset of patients, the use of previously annotated genetic information from re-analyzed studies^14,15,22^, and *de novo* profiling of in-house patient samples with single-cell targeted DNA sequencing (scDNA-seq) with immunophenotyping through the Tapestri platform. We used these profiles to identify genetic subclones within individual patients and evaluated whether their lineage priming differed (**Figure 5A**).

**Figure 5.**
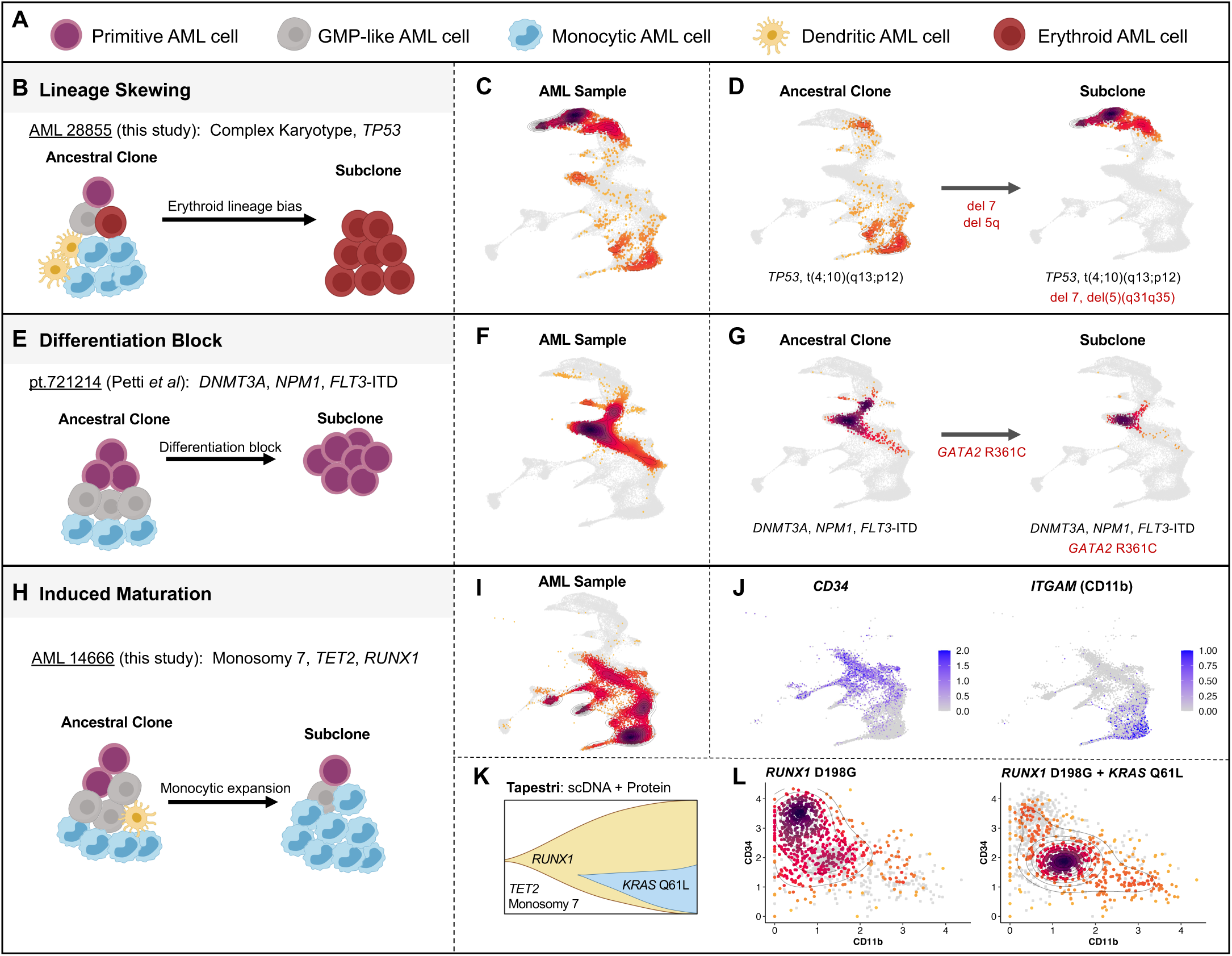
Impact of genetic subclones on AML differentiation. **A)** Graphical representation of broad differentiation states within AML samples. **B-D)** Example of lineage skewing from myeloid to erythroid at the sub-clonal level, depicting projection results from **C)** the primary AML sample and from **D)** an ancestral clone involving t(4;10) and an erythroid-restricted sub-clone involving t(4;10), del(5q), and monosomy 7. **E-G)** Example of a primitive differentiation block at the sub-clonal level, depicting projection results from **F)** the primary AML sample and from **G)** an ancestral *GATA2* wildtype clone and an HSC/MPP-restricted *GATA2-*mutant sub-clone. **H-L)** Example of induced maturation of AML cells at the sub-clonal level, depicting projection results from **I)** the primary AML sample, wherein **J)** primitive stem and progenitor cells express *CD34* and mature myeloid cells express *ITGAM* (encoding CD11b). **K)** scDNA and protein profiling of this sample by the Mission Bio Tapestri platform reveals an ancestral clone with monosomy 7 and *TET2* and *RUNX1* mutations with a sub-clonal *KRAS-*mutant clone. **L)** Immunophenotype analysis reveals the *RUNX1-*mutant ancestral clone to be mostly primitive (CD34+CD11b-) and the *KRAS*-mutant sub-clone to be mostly mature (CD34-CD11b+).

We found examples of subclonal mutations directly impacting differentiation landscapes in AML, including lineage skewing from myeloid to erythroid in a *TP53-*mutant subclone after deletion of chr7 and chr5q (**Figure 5B-D**), or differentiation block at the level of HSC/MPPs induced by a subclonal *GATA2* mutation (**Figure 5E-G**). In one sample that was profiled by scRNA-seq alongside Tapestri scDNA-seq with immunophenotyping (**Figure 5H)**, we identified CD34-expressing primitive cells co-existing with CD11b-expressing mature monocytic cells (**Figure 5I-J**). Through scDNA-seq with immunophenotyping from this same sample, we found that cells belonging to an ancestral Monosomy 7 and *RUNX1*-mutant clone were predominantly primitive (CD34+CD11b-), whereas cells that belonged to a *KRAS*-mutant subclone were predominantly mature (CD34-CD11b+) (**Figure 5K-L**). This approach enabled the functional annotation of subclonal genetic mutations with defined hematopoietic perturbations including lineage restriction, differentiation blocks, and even mature myeloid cell expansion within individual AML patients.

### Non-genetic heterogeneity within *KMT2A*-rearranged AML

Having mapped the associations between genetic heterogeneity and differentiation landscapes in AML, we next sought to explore non-genetic sources of heterogeneity between AML patients. Specifically, we considered heterogeneity in the origins of disease-propagating LSCs at the apex of these hierarchies. While LSCs retain a primitive identity in most patients^65,80–83^, phenotypically committed, non-quiescent LSCs have also been reported within a subset of AML patients usually containing *KMT2A* rearrangements or *NRAS/KRAS* mutations^84–87^ and in some cases these have been shown to co-exist^27,88,89^.

To investigate LSC heterogeneity in AML, we first turned to bulk transcriptomes from *KMT2A*-rearranged (*KMT2A*-r) AML patients, where committed LSCs have been most commonly observed^84–86^ and where both cellular and transcriptional heterogeneity has been reported^70,90^. Consensus clustering by AML differentiation stage abundance across 113 *KMT2A*-r patients revealed two subgroups of *KMT2A*-r AML, termed *KMT2A*-r Early (n=27) and *KMT2A*-r Committed (n=86) (**Figure 6A**). While multiple fusion partners were observed within each *KMT2A*-r subgroup, *KMT2A::MLLT4* was over-represented among *KMT2A*-r Early samples (*P=*4e-6) while *KMT2A*::*MLLT3* was over-represented among *KMT2A*-r Committed samples (*P=*0.007) (**Figure 6B**). *KMT2A*-r Early samples featured stem and progenitor involvement (HSC/MPP, LMPP, MEP) while *KMT2A*-r Committed samples were often entirely mature (ProMono, Monocyte, cDC) without involvement of stem and progenitor cells (**Figure 6C-D**). Strikingly, *KMT2A-*r Early samples were highly enriched for signatures of primitive LSC^65^ and quiescent LSPC^29^, while *KMT2A*-r Committed samples were highly enriched for signatures of committed LSC^86^ as well as cell cycle progression (**Figure 6E**), suggesting that these subgroups are maintained by distinct LSC populations. Thus, despite their shared genetic driver alterations, biological differences between these subgroups may reflect the distinct identities of their disease-propagating LSCs, either resembling primitive stem cells or committed myeloid precursors that have aberrantly acquired self-renewal capabilities (**Figure 6F**).

**Figure 6.**
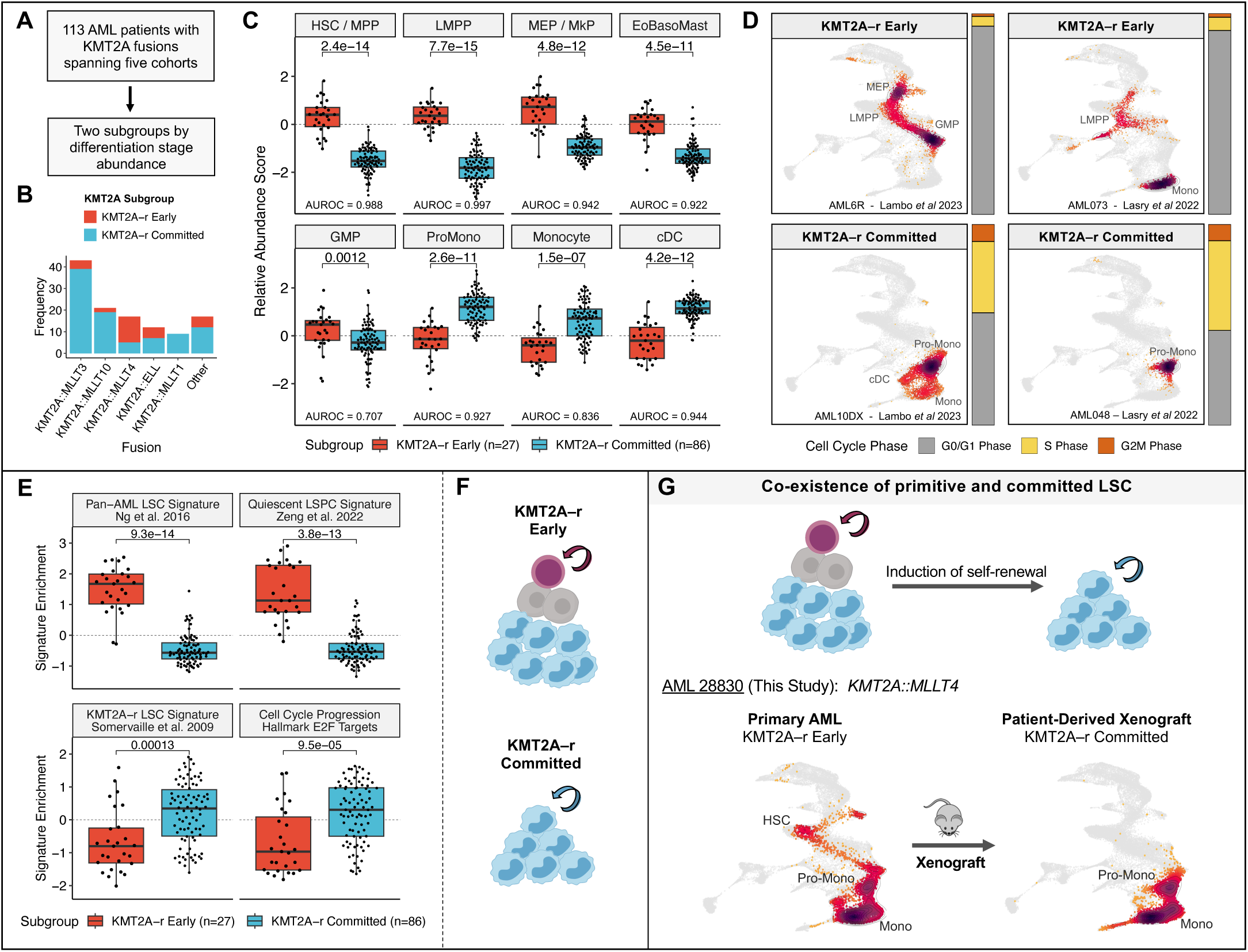
Non-genetic heterogeneity within KMT2A-rearranged AML. **A)** Identification of patient subgroups within 113 KMT2A-rearranged AML patients based on differentiation stage abundance. **B)** Distribution of KMT2A fusion partners based on KMT2A-r Early and Committed subgroups. **C)** Relative abundance of AML differentiation stages between KMT2A-r Early and Committed subgroups. P values by Wilcoxon rank-sum test along with the area under the receiver operating characteristic curve (AUROC) are shown for each comparison. **D)** Representative patient samples from the KMT2A-r Early and KMT2A-r Committed subgroups. The distribution of cell cycle phases is also shown for each sample. **E)** Gene set variation analysis (GSVA) enrichment of primitive LSC signatures against a KMT2A-r specific committed LSC signature, as well as E2F targets, across Early and Committed subgroups of KMT2A-r AML. **F)** Model for hierarchical organization of Early and Committed subgroups of KMT2A-r AML. **G)** Example of co-existing primitive and committed LSC within a KMT2A-r Early sample, wherein the patient-derived xenograft is independently sustained by a committed subclone independently.

Recognizing that *KMT2A*-r AML could be driven by primitive LSC or committed LSC, we asked whether these LSC populations could co-exist within the same patient. To evaluate this, we performed xenotransplantation with a *KMT2A*-r Early sample and mapped the single cell transcriptomes from the patient-derived xenograft (PDX). Strikingly, this PDX was engrafted by a fully committed clone, suggesting the pre-existence of a committed LSC in the primary sample (**Figure 6G**). Collectively, these findings nominate heterogeneity in LSC identity as a source of non-genetic heterogeneity within *KMT2A*-r AML and suggest that committed LSC may be formed by a direct transformation event or by evolution from a primitive LSC.

### Distinct leukemia cell hierarchies can co-exist within individual AML patients

To formally determine whether distinct LSC-driven hierarchies may co-exist within AML patients, we re-analyzed engraftment data from 74 AML patients wherein leukemia cells were sorted into four fractions based on surface expression of CD34 and CD38 and transplanted into NSG mice at varying doses^65^. Functional LSC activity, defined as the ability to initiate leukemic grafts, was observed within the CD34+CD38-fraction for 56 out of 74 patients (76%). Of the remaining 18 patients (18%), 7 patients (9%) had LSC activity detected within the CD34+CD38+ fraction whereas 11 patients (15%) had LSC activity restricted to the CD34-fraction. Considering the majority of patients with LSC activity within the CD34+CD38-fraction, 8 patients (11%) had LSC activity restricted to CD34+CD38-cells and 15 patients (20%) had LSC activity restricted to CD34+ fractions. These engraftment data are outlined in **supplemental Figure 12**.

Interestingly, 29 patients (39%) had detectable LSC activity in three or more fractions spanning CD34+ and CD34-immunophenotypes. The presence of LSC activity across multiple fractions could be related to previously documented immunophenotypic plasticity of CD34 and CD38 expression among primitive LSCs^65,82,83,91–94^, particularly in patients with primitive hierarchies with extensive stem and progenitor involvement^29^. In other patients, this pattern might instead be explained by the co-existence of leukemia cell hierarchies sustained by a combination of primitive and mature populations with functional LSC properties. To explore the latter possibility, we evaluated the immunophenotype of PDXs generated from each CD34 / CD38 fraction and identified two patient samples with discordant PDX immunophenotypes following transplantation of primary AML fractions sorted on CD34+ and CD34-for further analysis.

For these two patients, we performed scRNA-seq on the unsorted primary patient sample as well as sorted CD34+CD38-(34+38-) and CD34-CD38+ (34-38+) cell fractions. We transplanted these CD34+CD38- and CD34-CD38+ fractions separately and performed scRNA-seq on cells from the resulting PDX (**Figure 7A**). Flow cytometric and scRNA-seq analysis of primary cells from one patient (pt.90240) revealed multiple leukemia cell populations spanning LMPP-like, common lymphoid progenitor (CLP)-like, GMP-like, Pre-DC-like, and Mono-like stages (**Figure 7B-C, supplemental Figure 13**). Within the CD34+CD38-fraction (0.06% of AML cells), nearly all cells were LMPP-like (**Figure 7D**). PDX samples derived from this fraction recapitulated the complete patient hierarchy by both flow cytometric and scRNA-seq analysis (**Figure 7E**). The CD34-CD38+ fraction (11% of AML cells) comprised primarily downstream progenitors (**Figure 7F**). Critically, PDX samples derived from this fraction were almost entirely CD34- and composed of dendritic cell precursors and mature monocytes (**Figure 7G**). Similar results were observed with the second patient (pt.90394), wherein the CD34+CD38-graft recapitulated the full AML hierarchy while the CD34-CD38+ graft was composed entirely of mature myeloid cells (**supplemental Figures 14 and 15**). Collectively, these data provide proof of principle for a model wherein distinct leukemia cell hierarchies may co-exist within a rare subset of individual AML patients (**Figure 7H**), which may in turn contribute to AML evolution and relapse following selective pressures applied by different therapies (**Figure 7I**).

**Figure 7.**
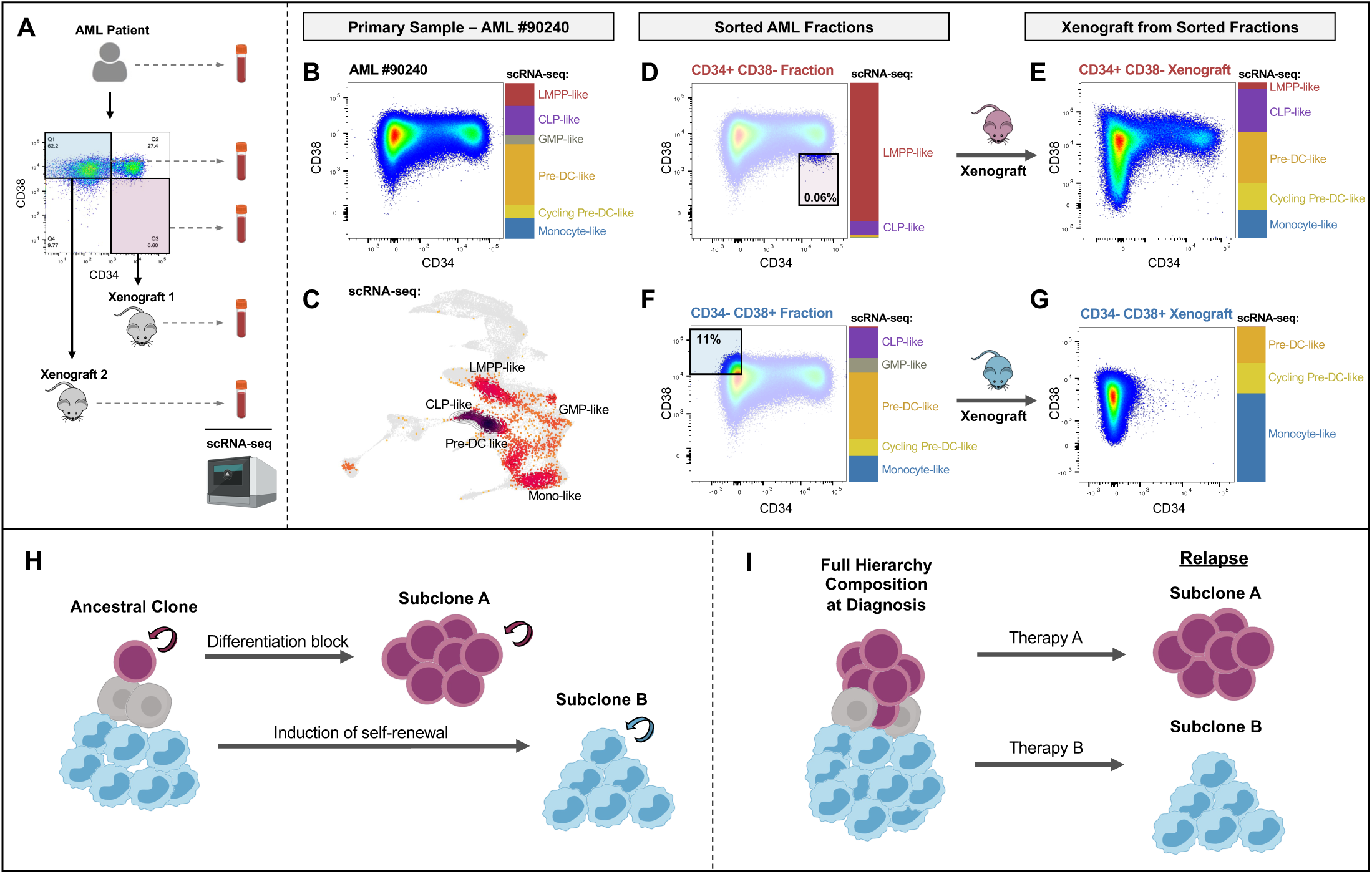
Distinct leukemia cell hierarchies can co-exist within individual AML patients. **A)** Outline of experimental workflow for identifying co-existing LSC-driven hierarchies in AML. Briefly, the primary AML patient sample, CD34+CD38-fraction, and CD34-CD38+ fraction are profiled by flow cytometry and scRNA-seq. Patient-derived xenografts (PDXs) from the CD34+CD38-fraction and from the CD34-CD38+ fraction are also profiled by flow cytometry and scRNA-seq and the composition of their leukemia cell hierarchies are compared. **B-G)** Results for AML patient #90240. **B)** Immunophenotype and scRNA-seq cell state composition of primary AML. **C)** scRNA-seq projection results of primary AML sample. **D)** Immunophenotype and scRNA-seq composition for CD34+CD38-fraction, representing 0.06% of primary cells. **E)** Immunophenotype and scRNA-seq composition for PDX derived from CD34+CD38-fraction, recapitulating the primary AML hierarchy. **F)** Immunophenotype and scRNA-seq composition for CD34-CD38+ fraction, representing 11% of primary cells. **G)** Immunophenotype and scRNA-seq composition for PDX derived from CD34-CD38+ fraction, restricted to mature AML populations. **H)** Model depicting the co-existence of leukemia cell hierarchies within a subset of AML patients, which could originate from intra-tumoral genetic heterogeneity. **I)** While relapse to conventional therapies is associated with expansion of primitive clones, acquisition of self-renewal at later stages of AML differentiation could enable expansion of self-sufficient monocytic clones at relapse to novel therapies targeting primitive AML cells.

## Discussion

Our reference map of bone marrow hematopoiesis provides in-depth characterization of human hematopoietic differentiation from early HSPC to terminally differentiated populations spanning multiple lineages. Most importantly, through careful annotation and validation, this curated map constitutes a valuable resource for those planning to utilize scRNA-seq data to study normal and malignant hematopoiesis. To maximize the public utility of our resource, we have developed an R package (https://github.com/andygxzeng/BoneMarrowMap) to enable rapid reference mapping and annotation of new scRNA-seq data across the spectrum of normal and malignant hematopoietic contexts.

The application of this reference mapping approach for classification of leukemia cells, particularly in AML, has allowed us to approach disease heterogeneity in a biologically meaningful way. Strikingly, some AML samples exhibited virtually no overlap in cell state involvement with one another, likely reflecting the impact of specific genetic drivers along with their disparate cellular origins. In contrast, other AML samples shared extensive overlap in cell state involvement with *bona fide* MPAL or AEL samples. These disease classes are discerned from AML by strict immunophenotypic criteria involving a handful of markers^7^, yet our evaluation of thousands of expression-based lineage markers suggests that additional markers may be needed to confidently identify malignant lymphoid and erythroid progenitor states. Translation of the new markers uncovered through scRNA-seq may permit the derivation of disease classes that are more homogeneous in their lineage identity, thereby enabling improved disease classification and therapeutic targeting.

Despite the various genetic pathways through which hematopoietic differentiation is disrupted in acute myeloid leukemia, these diverse perturbations converge upon twelve recurrent patterns of aberrant differentiation. We propose that each differentiation pattern reflects at least two key aspects of leukemogenesis. First, they reflect the identity of the last normal or pre-leukemic cell which transformed into leukemia, the cell-of-transformation^95^, which is likely mirrored by the LSC that sustains the disease. Importantly, the identity of this cell can differ from the cell-of-origin in which the first hit occurred. While this appears to be an HSC, MPP, or LMPP for most cases (p1-p4; p6-p9), in a minority of cases it appears to be a committed progenitor that had acquired self-renewal properties (p5; p10-12). Second, they reflect the available differentiation trajectories upon which the LSC can embark. These ‘rules’ of differentiation are determined by the genetic driver alteration. For example, in cases where the LSC identity mirrors that of a multipotent stem cell, differentiation may nonetheless be restricted to a single lineage due to the influence of their genetic driver on downstream lineage transitions. Moreover, depending on the genetic driver, differentiation may also be blocked at the stem cell stage (p1), blocked at the progenitor stage (p2-p4; p7), or fully permitted within a given trajectory (p8-p9). We propose that these two elements - the identity of the LSC and the ‘rules’ of downstream differentiation imposed by the genetic driver alteration - are central to understanding aberrant differentiation in AML.

This study also represents the most comprehensive effort to map genotype-to-phenotype associations between driver mutations and leukemic differentiation. Our findings align with existing knowledge from individual studies of genetic mutations, further validating the link between specific mutations and phenotypic changes in differentiation. By analyzing AML genetics at the single-cell level, we have built upon previous studies^2,3,22,96^ to show that distinct genetic subclones can differ in their cellular hierarchy composition, and formally demonstrated that distinct AML hierarchies sustained by distinct LSC populations can co-exist within a subset of patients. The co-existence of primitive and mature AML populations with functional LSC properties was previously proposed in a subset of AML patients^88,97^, with monocytic LSCs emerging as an important source of relapse from Venetoclax-based therapy^27,89^. Recognizing and accounting for this heterogeneity in LSC populations will be essential to improving the therapeutic targeting of AML.

## Acknowledgements

We thank the patients for sample donation. We thank Mason Boulanger for help with organization of clinical data and all members of the Dick lab for helpful discussions. A.G.X.Z is supported by a University of Toronto MD/PhD studentship award. J.E.D is supported by Princess Margaret Cancer Foundation; Ontario Institute for Cancer Research through funding provided by the Government of Ontario; Canadian Institutes for Health Research RN380110-409786; International Development Research Centre Ottawa Canada; Canadian Cancer Society 703212; Terry Fox New Frontiers Program project grant 1106; University of Toronto’s Medicine by Design initiative with funding from the Canada First Research Excellence Fund; The Ontario Ministry of Health; Canada Research Chair.. C.G.M is supported by the American Lebanese Syrian Associated Charities of St. Jude Children’s Research Hospital; the Alex’s Lemonade Stand Foundation for Childhood Cancer; the National Institutes of Health, National Cancer Institute grants P30 CA021765 and R35 CA197695; a St. Baldrick’s Foundation Robert J. Arceci Innovation Award; the Henry Schueler 41&9 Foundation.

## Authorship Contributions

A.G.X.Z.: designed and led the study, performed functional experiments, analyzed data, and wrote the manuscript. I.I.: performed single cell profiling experiments and analyzed data. S.S.: analyzed data and contributed to figure preparation. A.M.: performed functional experiments and contributed to engraftment analyses. S.B.: contributed to engraftment analyses and figure preparation. D.C.: analyzed data. G.W.: performed functional experiments. Q.G.: analyzed data. H.K.: contributed to functional experiments. J.A.K.: contributed to engraftment analysis. A.A. and M.D.M.: provided clinical samples. T.H.: provided clinical samples. C.G.M.: provided clinical samples. J.E.D.: contributed to study design, supervised the study, and revised the manuscript for final approval. All authors contributed to manuscript writing and gave final approval.

## Data and Code Availability

The BoneMarrowMap atlas can be explored interactively through the cellxgene web portal: https://cellxgene.cziscience.com/e/cd2f23c1-aef1-48ae-8eb4-0bcf124e567d.cxg/. An R package for projecting and classifying hematopoietic cells from query scRNA-seq data is available at (https://github.com/andygxzeng/BoneMarrowMap). For earlier access to datasets from this study, including processed single-cell transcriptomes from normal and leukemic samples, please contact Andy G.X. Zeng and John E. Dick.

