## Supplemental for "Single-cell transcriptional mapping reveals genetic and non-genetic determinants of aberrant differentiation in AML"

### Supplemental Note 1

#### Comparison of transcriptional cell states and functionally-defined HSPC subsets

To validate the reference map and cell state assignments, transcriptional cell state annotations were compared against functionally-defined HSPC populations. In addition to transcriptional projection of bulk transcriptomes from flow-sorted populations along the hematopoietic hierarchy [1–3] (**Figure 1B**), we utilized multiple additional datasets to benchmark our cell state annotations (**supplemental Table 4**).

Multipotent progenitors (MPPs), common megakaryocyte-erythrocyte progenitors (MEPs), and common myeloid progenitors (CMPs) have previously been shown to be admixed populations, separable into multilineage fractions (F1: CD71- BAH1-) and Mk/Ery committed fractions (F2: CD71+ BAH1-; F3: CD71+ BAH1+) [4]. Projection of transcriptomes from these subfractions showed that multi-lineage F1 fractions and Mk/Ery restricted F2 and F3 fractions within MPPs, CMPs, and MEPs were positioned in alignment with their known functional characteristics (**supplemental Figure 1A**).

Immunophenotypically-defined long-term (LT) HSC fractions exhibited high concordance with transcriptional cell state annotations (**Figure 1C, supplemental Figure 1C,E**). Notably, increasing concordance between transcriptionally- and immunophenotypically-defined HSCs was observed with increasing stringency of markers used in HSC purification[5–15], surpassing 90% concordance among LT-HSC fractions purified with additional markers including CD49f+[11,13], CD49f+CD34$^{lo}$CLEC9A$^{hi}$[14], and CD49f+EPCR+[15] (**supplemental Figure 1D**).

We next projected scRNA-seq profiles of hematopoietic progenitors including MPPs, lympho-myeloid primed progenitors (LMPPs), multi-lymphoid progenitors (MLPs), CMPs, MEPs, and granulocyte-monocyte progenitors (GMPs). Stringently-gated populations of LMPP, GMP, and MLP from Karamitros *et al* [16] were well-aligned with transcriptionally defined cell states by scRNA-seq projection (**supplemental Figure 1B)**. Yet, in other datasets immunophenotypic fractions including MPPs, lympho-myeloid primed progenitors (LMPPs), CMPs, MEPs, and granulocyte-monocyte progenitors (GMPs) were transcriptionally admixed and spanned multiple cell states [13] (**supplemental Figure 1F-L**), consistent with prior reports of functional heterogeneity within these fractions [4,16–18]. Indeed, re-analysis of index-sorted scRNA-seq profiles [10] of GMPs and MEPs revealed that CD38$^{mid}$ cells were enriched for true progenitors while CD38$^{high}$ cells were enriched for more committed states (**supplemental Figure 2A-C**). In keeping with this observation, we found that individual CD38$^{mid}$ GMPs and MEPs formed colonies more efficiently than CD38$^{high}$ GMPs and MEPs in single-cell functional assays (**supplemental Figure 2D-J**).

#### Supplemental Note 2

##### Quantification of differentiation stage abundance in bulk RNA-seq

Differentiation stages in AML, identified by composition analysis, each span multiple cell states along hematopoiesis (**supplemental Figure 10A**). The relative abundance of each differentiation stage within each patient is represented by a score derived from composition-based NMF analysis as described above. To infer this abundance in bulk RNA-seq data, we identified biologically relevant genes correlated to the abundance of each AML differentiation stage by integrating three approaches of feature selection.

**First**, we correlated the abundance of distinct differentiation stages with the expression of individual genes across patient-level pseudo-bulk profiles. To do so, we included scRNA-seq samples from studies with ≥ 5 patient samples [5,19–34] and created patient-level pseudo-bulk profiles for 302 AML/MPAL samples from these studies. Pseudo-bulk samples were logCPM normalized and a Pearson correlation test was performed between normalized gene expression and abundance of each AML differentiation stage across the 302 patient samples. Filtering at FDR < 0.05 resulted in thousands of significantly associated genes for each differentiation stage, thus further filtering was required. First, adaptive K1 thresholding, adapted from the AUCell R package, was performed within significantly positively correlated genes and within significantly negatively correlated genes.

Next, we identified differentially expressed genes across AML differentiation stages. Cells belonging to each differentiation stage within an AML patient were pooled into pseudo-bulk profiles. Only profiles comprising ≥ 50 cells were retained. One-vs-all differential expression was performed between each differentiation stage and all others using DESeq2 (v1.32.0), correcting for Patient ID as a covariate. Top gene expression markers for each AML differentiation stage are visualized in supplemental Figure 10B.

**Together,** for each AML differentiation stage only the genes that meet these following criteria were used for model training:

1. Positively or Negatively correlated with abundance of this AML differentiation stage at FDR < 0.05 and exceeding an adaptive positive or negative threshold

2. Significantly positively or negatively differentially expressed in this AML differentiation stage compared to others at FDR < 0.01.

Using genes that met this criteria, LASSO regression was performed on developmental state abundance using logCPM normalized gene expression data from pseudo-bulk profiles of 160 AML patient samples, using R package glmnet (v4.1.2). Briefly, to optimize model parameters 5-fold cross-validation was performed within each dataset with 10 repeats. Data was shuffled in between repeats, resulting in performance estimates for 50 random train/test splits. Pearson correlations across each train/test split was used as a measure of accuracy on unseen data (**supplemental Figure 10C**).

Final models for each developmental state were trained using LASSO, with leave-one-out cross-validation to determine the lambda value corresponding to the lowest root mean square prediction error (RMSE). To further reduce the number of features in the model, the largest lambda within one standard error of the lowest root mean square prediction error (RMSE) was used for the training of the final model. This resulted in a different gene expression model for inferring abundance of each AML differentiation stage (**supplemental Figure 10D-E**). All models are captured through a set of 400 genes (**supplemental Table 9**).

Thus, each score comprises a linear equation with coefficients/weights assigned by LASSO for each constituent gene. These equations are then applied to normalized bulk RNA-seq data from AML patients wherein normalized expression of each gene is multiplied by a corresponding coefficient, and the sum of these products is utilized and standardized across patients. Each standardized score represents the relative abundance of a developmental state within an individual patient. Scores are calculated in logCPM normalized gene expression data.

### Supplemental Methods

#### BoneMarrowMap Construction

##### Included Datasets

To develop a map with balanced representation of early hematopoietic stem and progenitor cells (HSPCs) alongside differentiated immune populations, we combined two bulk bone marrow datasets [35,36] with three CD34+ sorted bone marrow datasets [37–39] and a final dataset containing both bulk and CD34+ sorted samples [34], comprising six datasets and 45 unique donors. For the HCA Bone Marrow dataset, we utilized pre-filtered data from Hay *et al* [40]. Due to the high number of donors, variation in sorting approach, and complex relationships between cell populations along the continuous differentiation landscape, we found that restricting the inclusion criteria of datasets to 10x V2 chemistry and bone marrow tissue source helped to improve the quality of batch correction.

##### Preprocessing and Doublet filtering

scRNA-seq data from each study was preprocessed with scanpy [41] and Seurat [42], filtered at nGenes > 500 and pct.mito < 8%. To remove doublets from the dataset, we over-clustered the data and identified clusters with high proportions of doublets inferred through scrublet [43]. For this process, feature selection included the intersection of the top 3000 highly variable genes (HVGs) when stratifying by study and the top 3000 HVGs when stratifying by donor. This resulted in 2386 variable genes used for downstream dimensionality reduction. After principal component analysis (PCA), batch correction was applied across donors using harmony [44]. Leiden clustering [45] was performed using the top 50 harmony-corrected PCs and the top 50 nearest neighbours for each cell.

Doublet scores were calculated independently by applying scrublet to cells from each individual donor and the jenks natural breaks algorithm was applied to identify a universal doublet score threshold wherein cells with scores above this threshold were flagged as possible doublets. Each of the 64 clusters was evaluated to determine the proportion of cells above this cutoff, leading to the removal of nine clusters wherein >25% of the cells were flagged as potential doublets. We excluded these nine clusters from further analysis. This resulted in a total of 263,159 single cells used for generation of the hematopoietic reference map.

##### Dimensionality Reduction

Low-dimensional embeddings derived from scRNA-seq data are highly dependent on data pre-processing, variable gene selection, batch correction parameters, and dimensionality reduction-specific parameters. As a result, specific relationships between cell populations inferred through dimensionality reduction as well as downstream applications such as pseudo-time analysis will also vary depending on the parameters used.

To address this we performed a semi-supervised approach for choosing a final low-dimensional UMAP embedding based on prior biological knowledge. Specifically, we performed an iterative grid search of variable gene selection, batch correction, and UMAP parameters to identify an embedding that optimized specific constraints of hematopoietic differentiation based on prior biological knowledge.

Specifically, the following constraints were used:

- **Continuity of hematopoietic differentiation manifold.** With the exception of T/NK, Plasma, and Stromal populations, we required a continuous manifold capturing differentiation across the entire hematopoietic hierarchy. Embeddings with continuous two-dimensional manifolds were favoured.

- **Positioning of Pro-B and Pre-B cells** One important point of variability across UMAP embeddings pertained to the positioning of CD34+ Pro-B and CD34- Pre-B cells, and the transitional cycling state between these two stages of B cell development. Parameters that overemphasize cell cycle genes beyond B-cell development genes fail to position the transitional cycling state between Pro-B and Pre-B states. Embeddings that maintained a stepwise progression from Pro-B to Cycling to Pre-B were favoured.

- **Positioning of pDC development** The position of plasmacytoid DCs (pDCs) and their precursors relative to other populations also displayed variability across UMAP embeddings. Focused clustering of pDC precursors showed that they branched off from Multi-Lymphoid Precursors (MLPs) prior to restricted lymphoid development. We thus favoured embeddings that showed continuity between MLPs and Pre-pDCs.

- **Positioning of pDCs and cDC** pDCs and cDCs have been shown to be connected through a transitional AXL+ SICLE6+ "ASDC" state [46]. In addition to the prior constraint, we favoured embeddings that positioned pDCs proximally to cDCs through the ASDC population.

By adjusting the batch correction parameters (e.g. correcting on Study and Donor, or just correcting on Donor; adjusting theta for strength of batch correction; adjusting the tau parameter to avoid over clustering, and adjusting the "nclust" parameter to change the number of soft clusters harmony identifies in order to perform batch correction. UMAP parameters were also modified in addition to harmony parameters: these included the number of neighbours included in the neighbourhood graph (30 or 50) and the minimum distance between points (min_dist, spanning from 0.10 - 0.35). The resulting UMAP embeddings from each set of parameters were evaluated. Parameters for the final embedding included:

- 2386 highly variable genes. This is the intersection of the top 3000 highly variable genes when blocking by donor and the top 3000 highly variable genes when blocking by study.

- Harmony correction on two variables: "Study" with theta = 0, and "Donor" with theta = 1. Nclust = 70 and max.iter.cluster = 100.

- UMAP from 50 harmony components with k = 50 neighbours, min.dist = 0.23, spread = 1, and distance metric = 'cosine'.

#### Cell State Annotations

To derive cell state annotations, we performed graph-based leiden clustering across different resolutions (from 0.5 to 20). We found that unbiased metrics to select the optimal number of clusters such as the silhouette score were highly correlated with the number of clusters and inappropriate for use. Instead, we employed a flexible cluster assignment approach, wherein leiden clusters assigned at different resolutions were evaluated for their concordance with biological priors and assigned as specific cell states accordingly, with boundaries between cell states being informed by this prior information. These priors were dependent on the specific lineage being annotated:

- For differentiated cell populations including T cells, NK cells, B cells, Plasma cells, stromal cells, megakaryocytes, eosinophil/basophil/mast cells, pDCs, and cDCs, we leveraged existing annotations from bulk Bone Marrow references including Hay *et al*, Seurat Azimuth, Granja *et al*, and van Galen *et al* [5,34,40,42].
- For Hematopoietic stem and progenitor cell populations (HSPCs), we performed SingleR scoring from a bulk RNA-seq reference dataset from stringently purified HSPC populations [47] alongside AUCell [48] scoring of gene expression signatures of quiescent and activated HSCs [47,49]. HSPC population annotations were also refined based on external scRNA-seq validation datasets of sorted HSPCs and an in-house dataset of sorted immunophenotypic LT-HSCs.
- For populations along B cell development, we performed SingleR scoring from an in-house bulk RNA-seq reference dataset [50] from purified B cell precursors (CLP: CD34+CD38+CD10+CD19-; Pre-Pro-B: CD34+CD38+CD10-CD19+; Pro-B: CD34+CD38+CD10+CD19+; Pre-B: CD34-CD38+CD10+CD19+; Mature B) and utilized previously reported marker genes across stages of B cell development.
- For populations along erythroid development, we performed SingleR scoring from a bulk RNA-seq reference dataset [3] with purified erythroid populations (Pro-Erythroblast, Basophilic Erythroblast, Polychromatic Erythroblast, Orthochromatic Erythroblast), and utilized previously reported marker genes across stages of erythroid development.
- Established bulk RNA-seq profiles from sorted human populations between GMP and Monocytes or cDCs are not available, with the primary basis of these populations (e.g. early and late pro-monocytes) being morphological. Given the absence of clear ground truth reference profiles, we placed an emphasis on existing transcriptional annotations of myeloid development, evaluating the labels from Granja *et al* [34] and projecting labeled BM data from van Galen *et al* [5].
- As an additional layer of validation, whole transcriptome + Abseq single cell profiles from Triana *et al* [18] were projected onto the reference map to evaluate the concordance between assigned cell types and surface marker profiles.

#### BoneMarrowMap Transcriptional Characterization

##### Differentially Expressed Marker Genes

To find reliable markers for each cell state, a composite score was derived from seven distinct differential expression statistics, outlined below.

At the single cell level, the wilcoxauc function from presto was run to get the following statistics: 1) padj from Wilcoxon test on single cells, 2) log2 fold change (log2FC) from single cells in group vs single cells from all other groups, 3) AUC metric for distinguishing between cells in group vs cells from all other groups, 4) Log2 Detection Ratio between cells in group vs cells from all other groups. Including the AUC and Log2 Detection Ratio help to filter out normalization artifacts which are highly expressed at baseline, including ribosomal and mitochondrial genes.

Next, pseudo-bulk profiles were created by combining cells based on CellType & Donor (excluded pseudobulks with <5 cells) and DESeq2 was used to compare pseudobulks from each cell state against those from all other cell states, including “Study” and “Tissue” as covariates and evaluating significance through an LRT test. Based on this pseudo-bulk analysis, the following statistics were also incorporated: 5) padj from DESeq2, 6) log2FC from DESeq2. Pseudo-bulk profiles were subsequently *vst* normalized and wilcoxauc was run to produce 7) an AUC statistic at the pseudo-bulk level.

The geometric mean from all seven statistics was used to prioritize marker genes for each cell state, and subsequently called “MarkerScore”.

##### Consensus Non-negative Matrix Factorization:

Consensus non-negative matrix factorization (cNMF) [51] was first run with 10 iterations on the single cells from components 40-60 to help guide component selection by assessing the silhouette score (stability) and Frobenius reconstruction error (error). We used raw counts in the –counts flag and SCTransform V2 corrected counts from the data slot in the —tpm flag. After determining the optimal number of components (k=48), we ran 100 iterations of cNMF on the single cells while still inputting both the raw and SCTransform V2 non-negative corrected counts. The transcriptomic signatures were characterized using an overrepresentation analysis with the function, “fora” from the package fgsea [52] on the top 100 genes from each signature with HSC genesets and additional pathways from MSigDb.

##### pySCENIC Transcription Factor Regulon Inference

For transcription factor inference with SCENIC [48], the 263,159 single cells were aggregated to the metacell level using the divide and conquer algorithm from the Metacell2 package [53]. The highly variable genes used to construct the reference map were used as input into the algorithm guiding feature gene selection, thus optimizing clustering for metacell partitioning. The approach generated 10, 231 metacells with a target metacell size of 75,000 UMI. Gene regulatory network inference was performed using a candidate list of TFs [54], to identify co-expression modules using pySCENIC [55], particularly on the raw count metacells, to reduce runtime. Candidate regulons were pruned using the annotations of transcription factor motifs, 'motifs-v9-nr.hgnc-m0.001-o0.0.tbl.' Subsequently, Cistarget was employed using the 'mc9nr' databases, which include known human TF motifs annotated at: a) 500 bp upstream and 100 bp downstream of the transcriptional start site (TSS) and b) 10 kb centered around the TSS. Log-transformed counts of the Metacells were used as input for CisTarget, and drop-out masking was applied. Finally, the transcription factor regulons were scored using AUCell in the single-cell dataset, quantifying transcription factor activity.

##### Pseudotime Ordering

Monocle3 [56] was used to identify differentiation trajectories along the uniform manifold approximation and projection (UMAP) of the BoneMarrowMap, identifying 8 single-cell hematopoietic lineages of Erythroid, Megakaryocyte, EoBasoMast, Neutrophil, cDC, pDC, B-cell, and Monocyte development following prior biological knowledge. First, a principal graph was fitted to the UMAP using learn_graph with ncenter = 1200. For the pseudotime validation query data from Roy *et al* 2021 learn_graph was run with ncenter = 1400. Next, cells were ordered along the differentiation trajectories using order_cells from the "HSC" cluster, quantitatively characterizing development from primitive to mature states. The pseudotime ordering of the BoneMarrowMap was visualized across each trajectory using the package scPubr [57].

##### Transcription Factor Assignment Across cNMF Signatures

The inferred regulons from pySCENIC were used to construct a transcription factor gene set for overrepresentation analysis based on a hypergeometric test with the function "fora" from the fgsea package. Additionally, we computed Pearson correlation coefficients to compare the single cell cNMF usage scores and the transcription factor AUCell scores obtained from pySCENIC. We then calculated the geometric mean of the -log10(adjusted p-value) from "fora" and the Pearson single-cell correlation to derive a marker score for transcription factor prioritization. Transcription factors with an adjusted p-value greater than 0.01 and a pearson single-cell correlation below 0.3 were filtered out.

#### BoneMarrowMap Sample Projection

##### Projection of Normal and Leukemic Hematopoietic Samples

Single cell transcriptomes from normal and leukemic hematopoietic samples (query data) were projected onto BoneMarrowMap using Symphony [58] based on expression of 2386 variable genes defined from the reference map. Quality control is performed using a mapping error score adapted from Kang et al 2021 [58], represented by the mahalanobis distance of each query cell from the nearest centroid within the reference map. Filtering criteria are adapted to each dataset based on the distribution of the mapping error scores, with a threshold typically set between 1.5 and 3 median absolute deviations (MADs) above the median mapping error, wherein cells surpassing this mapping error threshold are excluded from downstream analysis.

Query cells are assigned to cell states by K-Nearest Neighbours (KNN) classification based on the identity of the 30 nearest neighbours from the reference map. After a cell state label is assigned, the confidence of this assignment is represented by the proportion of nearest neighbours from the reference map sharing this same label, hereafter “KNN probability”. This metric is used for filtering wherein confidently classified cells are retained for composition analysis, typically at a KNN probability > 0.5. Hematopoietic pseudo-time values were also predicted among query cells by KNN regression using the 30 nearest neighbours from the reference map.

#### Single-Cell Colony Assays

##### Human Cord Blood Samples

Human CB samples were obtained with informed consent from Trillium Health, Credit Valley and William Osler Hospitals according to procedures approved by the University Health Network (UHN) Research Ethics Board. Mononuclear cells were obtained by centrifugation on Lymphoprep medium (Stem Cell Technologies) and were depleted of Lin+ cells (lineage depletion) by negative selection with the StemSep Human Progenitor Cell Enrichment Kit according to the manufacturer’s protocol (Stem Cell Technologies). Lin- CB cells were stored viably at −80°C or −150°C.

##### Single Cell Colony Assays

For single cell colony assays, Lin- human CB cells were thawed via slow drop-wise addition of X-VIVO 10 medium with 50% fetal bovine serum (FBS) and DNaseI (200 µg/ml). Cells were spun at 350g for 10 minutes (min) at 4 °C and then resuspended in phosphate-buffered saline (PBS) + 2.5% FBS. From Lin- human CB cells, GMPs and MEPs were index-sorted with a gating strategy described in Garcia-Prat *et al* [47] and shown in **supplemental Figure 2D-F**. The following antibodies were used: CD34 APC7, CD38 PE-Cy7, CD90 APC, CD49f PE-Cy5, CD45RA FITC, CD10 AF700, CD19 AF700, CD7 AF700, FLT3 Biotin + Streptavidin Qdot PE, CD71 BV786. Cells were resuspended in 100 µl per 1x106 cells and stained in two subsequent rounds for 15 min at room temperature each.

Single cell colony assays were performed as previously described [59]. Three days prior to the single cell assay, Nunc 96-well flat bottom plates (Thermo Fisher) were treated with 50 µl 0.2 % gelatin solution (Sigma-Aldrich) per well for one hour. After the gelatin solution was removed, MS-5 murine stromal cells (*76*) were seeded at a density of 1,500 cells per well in 100 µl H5100 medium (Stem Cell Technologies). One day prior to the single cell assay, the H5100 medium was replaced with 100 µl of erythro-myeloid-megakaryocytic promoting medium, which consisted of StemPro-34 SFM medium (Thermo Fisher) with the provided supplement, 1x L-glutamine (Thermo Fisher), 1x penicillin-streptomycin (Thermo Fisher), 0.02 % human LDL (Stem Cell Technologies) and the following cytokines (Miltenyi Biotec unless stated otherwise): FLT3L (20 ng/mL), GM-CSF (20 ng/mL), SCF (100 ng/mL), TPO (100 ng/mL), EPO (3 ng/mL, Eprex), IL-2 (10 ng/mL), IL-3 (10 ng/mL), IL-6 (50 ng/mL), IL-7 (20 ng/mL) and IL-11 (50 ng/mL). On the day of the single cell assay, sorted single cells were deposited into the MS-5 seeded 96-well plates (80 wells/96-well plate) using the FACSAria II (BD). After 15-17 days, colonies from each single cell were scored after 15-17 days under the microscope.

#### AML scRNA-seq Profiling

##### MLL-SJCRH scAML - Single-cell RNA sequencing

Frozen mononuclear cells from AML samples from the Munich Leukemia Laboratories (MLL) were thawed, counted and enriched for live cells by depletion of dead cells (Miltenyi Biotec, catalog number 130-090-101). Isolated live cells were washed three times with 1X Phosphate-Buffered Saline (PBS, calcium and magnesium free) containing 0.04% weight/volume BSA (Thermo Fisher Scientific, catalog number AM2616) and automatedly counted by the Countess 3 Automated Cell Counter (Thermo Fisher Scientific) and manually by the Neubauer hemocytometer.  From each sample, we calculated the volume of cells to load to have a desired recovery target of 8,000-10,000 cells and loaded on Chromium Next GEM Chip G (10X Genomics; PN-2000177) with reagents from Chromium Next GEM Single Cell 5' Library & Gel Bead Kit v1.1 (10X Genomics; PN-1000165), following standard manufacturer's protocols for gel beads-in-emulsion (GEMs) generation, barcoding, cDNA amplification and gene expression library construction. Final library quality was assessed using High Sensitivity D1000 ScreenTape (5067- 5584) with High Sensitivity D1000 Reagents (5067- 5585) at the Agilent 2200 TapeStation system. Illumina-ready dual index libraries were sequenced at the recommended depth and aiming at 50,000 reads/cell at the Hartwell Center at St Jude Children’s Research Hospital (SJCRH) on the Illumina NovaSeq according to manufacturer's recommendations.

##### MLL-SJCRH scAML - Pre-Processing and Mapping

MLL-SJCRH scAML data were aligned and quantified using the Cell Ranger (v5.0.1) pipeline against genome GRCh38. Cells were filtered with the following criteria: nCount_RNA > 1000, nFeature_RNA > 500, pct.mito < 10), leaving a total of 116,839 primary AML cells profiled by 10x 5’ scRNA-seq. These cells projected onto BoneMarrowMap and filtered at a mapping error threshold of 2.5 MADs above the median.

##### Inference of Genetic Alterations from scRNA-seq

Inference of copy number variations (CNVs) was performed through inferCNV (https://github.com/broadinstitute/inferCNV) with the following parameters (cutoff = 0.1, HMM = TRUE, hmm_type = ‘i6’, BayesMaxPNormal = 0.2, analysis_mode = ‘subclusters’, cluster_by_groups = FALSE, tumor_subcluster_pval = 0.01, tumor_subcluster_partition_method = ‘qnorm’, denoise = TRUE). For the St. Jude scAML data, two CD34+ cord blood samples profiled by 10x 5’ scRNA-seq alongside the AML samples were used as CNV-neutral references. For the PMH scAML data, two mobilized peripheral blood samples were used as CNV-neutral references. When possible, called CNVs were validated with known CNV changes by WGS or karyotyping and validated CNV calls were used for downstream analysis. Expression mutations within scRNA-seq data were called using the cb_sniffer tool (https://github.com/genome/cb_sniffer) based on known genetic mutations from prior targeted sequencing of each AML sample, in line with Petti *et al* [33].

##### Targeted Single-cell DNA and Protein Sequencing

Single-cell amplicon-based DNA and protein sequencing was performed by using the Tapestri platform (Mission Bio), and a custom panel (Cell DNA Custom V2 Kit 2 MB03-0007 Panel ID: CO-131 and Supplementary Table 13), which assesses common AML mutations across 150 genomic regions of 36 genes. Eight oligo-conjugated antibodies from Mission Bio were used, including: anti-human CD45, anti-human CD45RA, anti-human CD34, anti-human CD38, anti-human CD33, anti-human CD90, anti-human CD11B, and anti-human CD123. Cryopreserved mononuclear cells were thawed, deprived of dead cells by using the Dead Cell Removal Kit (Miltenyi Biotech, #130-090-101), counted by the Countess 3 Automated Cell Counter (Thermo Fisher Scientific) and manually by the Neubauer hemocytometer. Cells were stained and processed following the Tapestri® Single-Cell DNA + Protein Sequencing User Guide standard manufacturer's protocols and the following kits from Mission Bio: Tapestri Single-Cell DNA Core (catalog number MB51-0007), Tapestri Single-Cell DNA Core (catalog number MB51-0010), Tapestri Single-Cell DNA Bead Kit (catalog number MB51-0009) and the Tapestri Custom Proteogenomic Panel. All libraries, both DNA and protein, were purified by Ampure XP beads, quantified and pooled for sequencing on an Illumina NovaSeq at the Hartwell Center at SJCRH according to manufacturer's recommendations.

##### Targeted Single-cell DNA and Protein Sequencing: analysis

FASTQ files for single-cell DNA and protein libraries were analyzed through the Tapestri pipeline in the Tapestri Portal. This pipeline trims adaptor sequences, detects barcodes, aligns reads to the human genome (hg19), assigns sequence reads to cell barcodes, and performs genotype calling. At the end of these steps, a VCF file is generated for each sample. This file is then used for the .loom file generation. Integrated DNA and protein analysis was performed by Mosaic v3.0.1 (https://github.com/MissionBio/mosaic), and surface protein abundance was subject to centered log-ratio normalization (CLR) for downstream analyses.

#### Single Cell AML Composition Analysis

##### Literature scAML - Pre-Processing and Mapping

scRNA-seq datasets from the literature were obtained and mapped onto BoneMarrowMap. When possible, pre-processed data from the original publication was used, otherwise custom QC filters were set based on the distribution of QC metrics (percent mitochondrial genes, number of unique genes detected in each cell) of each sample in a dataset. Custom mapping error filters were also applied based on the mapping error distribution for each sample within a dataset. These metrics vary from dataset to dataset depending on sample quality, library preparation technologies, and sequencing depth; they are highlighted in Supplementary Table 4.

##### Literature scAML - Composition Analysis

Combining our internal AML scRNA-seq dataset with eleven additional AML/MPAL datasets, we retained cells that passed mapping error QC and that were classified at a KNN probability ≥ 0.5.

After removing cells from healthy controls, patient-derived xenografts, and remission samples, this resulted in a total of 1,223,411 cells spanning 318 diagnosis or relapse samples from 274 unique patients (257 AML patients and 17 MPAL patients). For composition analysis of AML differentiation hierarchies, cells assigned to mature lymphoid (NK, T, B, Plasma) and Stromal states were removed, leaving 1,046,789 single-cell transcriptomes for composition analysis.

For composition analysis, cell states with less than 100 cells present across all samples were filtered out and the frequency statistics for each remaining cell state within each sample were used. Cell counts across these 37 cell states across 318 samples were centered log-ratio (CLR) normalized with multiplicative replacement using the package scikit-bio (v0.5.6). To simplify downstream analyses, we grouped specific cell states into broader categories based on correlated abundance across patient samples. To do so, we ran non-negative matrix factorization (NMF) on cell composition data with relative abundance of each cell type serving as the features, leading to the identification of multiple NMF components, each representing a group of correlated cell states. Thirteen NMF components were identified, each corresponding to discrete stages of differentiation along distinct branches of hematopoietic development, hereafter referred to as differentiation stages. To benchmark the abundance estimates for each differentiation stage, we compared the abundance across all cells versus their abundance across cells annotated as malignant by studies that performed a dedicated normal vs malignant analysis. The abundance of all differentiation stages was concordant with abundance estimates from malignant-only analyses ($r > 0.8$), with the exception of Late Erythroid ($r = 0.7$). Thus, abundances of the twelve differentiation stages (excluding Late Erythroid) across all cells within each patient sample were used for downstream composition analysis.

Differentiation stage abundance (derived from NMF) was standardized through z-scaling and used for consensus clustering through ConsensusClusterPlus v1.50.0 using pam and k-means algorithms separately, based on euclidean distances with 1000 repetitions up to a max K of 25. For each number of clusters K, silhouette scores were calculated based on (1) the NMF-derived matrix of 12 differentiation stages and (2) the CLR-normalized composition data of 37 cell states and the K values with the highest silhouette scores were retained for downstream analysis. Composition cluster assignments from these highest-scoring cluster assignments were integrated in a supervised manner through examination of cases with discordant assignments. This process yielded twelve patient subgroups, referred to as Differentiation Patterns, based on cellular composition differences along human hematopoiesis.

#### Lineage Quantification in Bulk RNA-seq

##### Quantification of differentiation stage abundance

Relative abundance of each differentiation stage was quantified as outlined in Supplemental Note 2 were calculated on logCPM-normalized bulk RNA-seq profiles from AML cohorts [60–65] and AEL cohorts[66]. For meta-analysis of AML cohorts, logCPM normalization was performed on COMBAT-seq batch corrected gene expression counts provided by Severens *et al* [65].

##### Statistical analysis in bulk RNA-seq cohorts

Unless otherwise indicated, all comparisons between groups were performed through a Wilcoxon rank sum test. Associations between differentiation stage abundance and cytogenetic characteristics were evaluated by a generalized linear model using glmnet (v4.1.3) wherein inferred abundance of each developmental state was used as the dependent variable and cytogenetic alteration status was used as the independent variable, stratified by patient cohort. This same approach was used for evaluating associations with mutation status, FAB classification, and hierarchy subtype. The resulting t-statistic and FDR corrected significance was visualized through package corrplot (v0.84).

##### Composition-based subclustering of KMT2A samples

113 *KMT2A*-rearranged (*KMT2A*-r) samples were identified across five AML cohorts based included in a meta-analysis from Severens *et al* [60–65]. To identify composition-based subclusters among these samples, consensus clustering was performed based on AML differentiation stage abundance using ConsensusClusterPlus v1.50.00 with the pam algorithm based on euclidean distance, with 1000 repetitions. Two *KMT2A*-r subclusters were identified that differed principally on whether stem and progenitor populations were present or absent. Published gene expression programs were calculated on logCPM normalized data using GSVA (v1.40.1).

#### AML Engraftment and LSC scRNA-seq Analysis

##### Primary AML sample engraftment analysis

AML cells from 74 primary patient samples were sorted into four fractions based on CD34 and CD38 expression and each fraction was transplanted into NSG mice at varying cell doses. Injected right femur engraftment at 12-weeks post-transplant was evaluated by flow cytometry. This data was reported in Ng *et al* [67]. Graft immunophenotype was used to assign AML fractions as either LSC+, LSC-, or multilineage. Fractions that initiated CD45+CD33+ leukemic grafts were denoted as LSC+ (dark blue), those that did not initiate grafts were denoted as LSC- (light grey), and those that initiated grafts with both CD33+ myeloid and CD19+ lymphoid cells were denoted as multilineage (green) and presumed to be either normal or pre-leukemic. Patient samples were categorized based on engraftment patterns across their CD34 and CD38 fractions to identify subsets with distinct patterns of AML engraftment.

##### Co-existing LSC: flow sorting, xenografting, and scRNA-seq library preparation

Primary AML samples from patients 90240 and 90394 were obtained from the Princess Margaret Hospital (PMH) biobank and subject to Fluorescence-Activated Cell Sorting (FACS) with the following panel: CD34-APC7, CD38-PE, CD33-BV786, CD45-APC, CD3-FITC. Using propidium iodide (PI) as a viability stain, the following live cell fractions were collected: Bulk AML (CD45+CD3-), CD34pCD38m (CD45+CD3-CD34+CD38-), CD34mCD38p (CD45+CD3-CD34-CD38+). For each collected cell fraction, cells from patients 90240 and 90394 were pooled together at similar proportions and subject to scRNA-seq with the 10x Next GEM Single Cell 3’ Gene Expression v3.1 kit and NovaSeq 6000 at a target depth of 50,000 reads per cell.

Sorted CD34pCD38m (CD45+CD3-CD34+CD38-) and CD34mCD38p (CD45+CD3-CD34-CD38+) fractions from primary AML patients 90240 and 90394 were transplanted into NSG mice at varying cell doses, with human leukemic engraftment in the injected right femur evaluated at 12 weeks post-transplant. The immunophenotype of the leukemic graft was evaluated with the following panel: CD34-APC7, CD38-PE, CD33-BV786, CD45-APC, CD3-FITC. Using PI as a viability stain, live CD45+ human leukemic cells were collected by FACS. CD34pCD38m-engrafted mice from patients 90240 and 90394 were pooled together at similar proportions and subject to 3' scRNA-seq as described above. CD34mCD38p-engrafted mice from patients 90240 and 90394 were also pooled together at similar proportions and subject to 3' scRNA-seq as described above.

As a non-leukemic control, two allogeneic mobilized peripheral blood (allo-mPB) samples were obtained from the PMH biobank. After thawing, CD34+ cells were purified using the CD34 Microbead UltraPure positive selection kit from Miltenyi. CD34- flowthrough cells were subject to dead cell removal using the EasySep Dead Cell Removal (Annexin V) kit from STEMCELL Technologies. Purified CD34+ and live CD34- cells from each donor were subsequently mixed at a 60/40 ratio of CD34+/CD34- cells. Mixtures from the two donors were subsequently pooled together at similar proportions and subject to 3' scRNA-seq as described above.

##### Co-existing LSC: scRNA-seq pre-processing and mapping

PMH scAML data were aligned and quantified using the Cell Ranger (v5.0.1) pipeline against genome GRCh38. Cells were filtered with the following criteria: nCount_RNA > 2000, nFeature_RNA > 1000, pct.mito < 15). Given that each scRNA-seq library consisted of two samples (one male and one female), SouporCell [68] was applied with k = 2 and default parameters to deconvolute scRNA-seq libraries by donor genotype and identify genotype-based doublets. Simultaneously, transcriptional doublets were identified using scDblFinder (v1.6.0) [69]. We validated the identity of the two donors identified by SouporCell based on expression scores for male-specific genes (ChrY, excluding the para-autologous region) and female-specific genes (XIST and TSIX). We validated that genotypic doublets by SouporCell and transcriptional doublets by scDblFinder exhibited co-expression of male-specific and female-specific genes; these doublets were filtered out. We further identified that <1% of single cells classified as male by SouporCell exhibited high levels of XIST and TSIX expression; these cells were filtered out.

After this QC and doublet filtering, high quality single-cell transcriptomes from 23,522 AML cells and 6383 allo-mPB cells were retained. These cells were projected onto BoneMarrowMap and filtered at a mapping error threshold of 5 MADs above the median. InferCNV analysis was performed as outlined above and compared against known clinical karyotyping profiles from AML90394 and AML90240 to distinguish leukemic cells from normal cells. To define leukemic sub-populations, clustering was performed separately within primary cells within each donor and within xenografted cells within each donor. Briefly, 20 or 30 principal components was used depending on the elbow plot for each sample, UMAP was run with n.neighbours = 30 and min.dist = 0.2, and louvain clustering with multilevel refinement was performed with res = 0.8, and AML subclusters were annotated based on assigned cell states from reference map projection.

#### Supplemental Table Legends

**Supplemental Table 1. Hematopoietic cell states in BoneMarrowMap**
List of 55 transcriptional cell states identified in BoneMarrowMap. Full names without abbreviations are also provided for clarity.

**Supplemental Table 2. Top marker genes for each hematopoietic cell state**
List of top 50 marker genes, ranked by differential expression (DE) enrichment, identified for each of the 55 cell states in BoneMarrowMap. DE was performed with DESeq2 using pseudobulk profiles from each cell state within each donor, with Donor and Study as covariates.

**Supplemental Table 3. Hematopoietic gene expression programs identified by cNMF**
Overview of 48 gene expression programs identified through unsupervised consensus NMF. The top 100 genes are shown for each program, ranked by NMF coefficient.

**Supplemental Table 4. Normal and Leukemia scRNA-seq datasets analyzed in this study**
Overview of published normal and leukemia scRNA-seq datasets re-analyzed in this study, with additional information pertaining to the contents of each dataset along with appropriate citations.

**Supplemental Table 5. Patient annotations from AML scRNA-seq meta-analysis**
Clinical and genomic annotations for 318 primary patient samples used for composition analysis from the MLL-SJCRH cohort and 20 additional scRNA-seq studies involving AML and MPAL patients. Standardized abundance scores for each AML differentiation stage are also shown, along with classification of each sample to one of twelve recurrent differentiation patterns.

**Supplemental Table 6. Cell state composition across 318 AML and MPAL patient samples**
Composition data for 318 patient samples across 38 cell states (excluding mature lymphoid and stromal cells), depicted as the number of cells mapping to each state in each patient sample.

**Supplemental Table 7. Differentiation stage definitions in AML composition analysis**
Consolidation of correlated AML cell states into broader differentiation stages for composition analysis. This is a list of the precise cell states underlying each AML differentiation stage.

**Supplemental Table 8. Marker Genes for each AML differentiation stage**
List of top 100 marker genes, ranked by differential expression (DE) enrichment, identified for each of the 13 Differentiation Stages across 318 AML and MPAL patient samples. DE was performed using pseudobulk profiles from each differentiation stage within each patient sample, with Study as a covariate.

**Supplemental Table 9. Models for predicting AML differentiation stage abundance**
Linear equations for predicting AML Differentiation Stage abundances in normalized bulk RNA-seq data (e.g. logCPM) from AML patients using a total of 400 genes.

**Supplemental Table 10. AML meta-analysis annotations and differentiation stage scores**
Quantified AML Differentiation Stage abundance across five AML cohorts (TCGA, BeatAML2, Leucegene, TARGET, and LUMC) totalling 1,224 patients. Clinical annotations provided by Severens *et al* 2024 were updated and hierarchy-based KMT2A-r subgroups are also included.

**Supplemental Table 11. Genotype-to-phenotype mapping across AML patients**
Results from genotype-to-phenotype association analysis between genetic driver alterations and AML differentiation stage abundance from the five-cohort dataset (n=1,224 patients). Statistics and p-values were calculated using a generalized linear model adjusting for the originating cohort as a covariate. Corrected q-values were obtained by the Benjamini-Hochberg method.

**Supplemental Table 12. AEL cohort annotations and differentiation stage scores**
Quantified Differentiation Stage abundance across 136 acute erythroid leukemia patients from Iacobucci *et al* 2019, along with clinical annotations provided by the original study.

**Supplemental Table 13. Custom Tapestri Panel**
Single-cell genotyping panel used for the Tapestri genotype and immunophenotype analysis of patient samples from the MLL-SJCRH cohort.

**Supplemental Table 14. Clinical annotations for the co-existing LSC patient samples**
Clinical and genomic annotations for two AML patient samples from Princess Margaret Hospital used for sorting, xenografting, and scRNA-seq profiling as part of the co-existing LSC analysis.

#### Supplemental Figures

<u>Supplemental Figure Headings:</u>

S1. Projection of sorted HSPC populations onto BoneMarrowMap

S2. Functionally distinct CD38 fractions within human GMPs and MEPs

S3. Identification of gene expression programs through consensus NMF

S4. Pseudotime analysis and characterization of erythroid and monocytic differentiation

S5. Characterization of B-lymphoid and dendritic cell differentiation

S6. Characterization of megakaryocyte and granulocyte differentiation

S7. Characterization of broad differentiation stages implicated in AML

S8. Expression of therapeutic targets and LSC signatures by AML differentiation stage

S9. Representative samples from each AML differentiation pattern

S10. Inference of AML differentiation stage abundance from bulk RNA-seq profiles

S11. Clinical and genomic correlates of AML differentiation stage abundance

S12. Analysis of engraftment patterns among CD34 and CD38 sorted fractions in AML

S13. Cell state classification in AML 90240 primary and xenograft samples

S14. Distinct LSC-driven leukemia cell hierarchies co-existing within AML 90394

S15. Cell state classification in AML 90394 primary and xenograft samples

**Supplemental Fig 1. Projection of sorted HSPC populations onto BoneMarrowMap**
**A)** Projection of purified multilineage (F1) and Mk/Ery restricted (F2, F3) sub-fractions of MPP, CMP, and MEP from Notta *et al* 2016. **B)** Projection of scRNA-seq profiles purified with functionally informed gating of LMPP, GMP, and MLP populations from Karamitros *et al* 2018. **C)** Projection of scRNA-seq profiles from HSCs purified using different markers of human HSCs. **D)** Proportion of transcriptional HSCs and MPPs projected from scRNA-seq data sorted with varying levels of immunophenotypic purity. The combination of markers used, tissue source, and dataset name are also depicted. **E-L)** Projection of scRNA-seq profiles of immunophenotypically defined HSPC populations from Zhang *et al* 2022, spanning **E)** HSC, **F)** MPP, **G)** LMPP, **H)** MLPs, **I)** Pre-B/NK, **J)** MEP, **K)** CMP, and **L)** GMP.

**Supplemental Fig 2. Functionally distinct CD38 fractions within human GMPs and MEPs**
**A-C)** Re-analysis of index-sorted scRNA-seq profiles from Velten *et al.* **A)** Gating of MEP, CMP, and GMPs within CD34+CD38+CD10- progenitors. **B)** Gating and scRNA-seq projection of CD38-mid and CD38-high GMP fractions. **C)** Gating and scRNA-seq projection of CD38-mid and CD38-high MEP fractions. **D-F)** Single-cell colony assays with index-sorting of CD38-mid and CD38-high **E)** GMP fractions and **F)** MEP fractions from umbilical cord blood. **G)** Single-cell colony formation outcomes within CD38-mid and CD38-high GMPs and **H)** surface CD38 levels of individual GMPs that did and did not form colonies. **I)** Single-cell colony formation outcomes within CD38-mid and CD38-high MEPs and **J)** surface CD38 levels of individual MEPs that did and did not form colonies.

**Supplemental Fig 3. Identification of gene expression programs through consensus NMF**
**A)** Unsupervised discovery of 48 gene expression programs (GEP) across human hematopoiesis by cNMF. 36 GEPs correspond to hematopoietic cell states (cell state programs) and 12 GEPs correspond to other cellular processes (activity programs). The relative activity of each program within each hematopoietic cell state is depicted through a heatmap of AUC values. **B)** Transcription factor (TF) regulons associated with each GEP based on a composite score representing both the correlation of GEPs with each TF regulon score as well as the significance of overlap between genes defining each GEP and each TF regulon. **C)** Key pathways significantly associated with activity GEPs. **D-G)** Cell cycle plot of S vs G2M scores within HSPCs, depicting **D)** cell cycle assignments and **E)** S-phase, **F)** G2-phase, and **G)** M-phase program expression. **H-I)** UMAP of hematopoietic cell states depicting **H)** cell cycle assignments and **I)** S-phase, **J)** G2-phase, and **K)** M-phase program expression.

**Supplemental Fig 4. Pseudotime analysis and characterization of erythroid and monocytic differentiation**
**A-C)** Projection and validation of hematopoietic pseudotime values in query data from Roy *et al* 2021. **A)** Pseudotime values calculated *de novo* on HSPC transcriptomes from Roy *et al* 2021 using Monocle3. **B)** Pseudotime values inferred through BoneMarrowMap projection of HSPC transcriptomes from Roy *et al* 2021. **C)** Comparison of *de novo* and projected pseudotime values across cell states annotated in Roy *et al* 2021. Average values for each cell state are depicted and the size and color of each cell state correspond to the number of cells within that state. **D-F)** Characterization of erythroid differentiation. **D)** Pseudotime score along erythroid

differentiation. **E)** Transcription factor regulon activity along erythroid differentiation pseudotime. **F)** Gene expression changes along erythroid differentiation pseudotime. **G-I)** Characterization of myeloid differentiation. **G)** Pseudotime score along monocyte differentiation. **H)** Transcription factor regulon activity along monocyte differentiation pseudotime. **I)** Gene expression changes along monocyte differentiation pseudotime.

**Supplemental Fig 5. Characterization of B-lymphoid and dendritic cell differentiation**
**A-C)** Characterization of B cell differentiation. **A)** Pseudotime score along B cell differentiation. **B)** Transcription factor regulon activity along B cell differentiation pseudotime. **C)** Gene expression changes along B cell differentiation pseudotime. **D-F)** Characterization of plasmacytoid dendritic cell (pDC) differentiation. **D)** Pseudotime score along pDC differentiation. **E)** Transcription factor regulon activity along pDC differentiation pseudotime. **(F)** Gene expression changes along pDC differentiation pseudotime. **G-I)** Characterization of conventional dendritic cell (cDC) differentiation. **G)** Pseudotime score along cDC differentiation. **H)** Transcription factor regulon activity along cDC differentiation pseudotime **(I)** Gene expression changes along cDC differentiation pseudotime.

**Supplemental Fig 6. Characterization of megakaryocyte and granulocyte differentiation**
**A-B)** Characterization of megakaryocyte differentiation. **A)** Pseudotime score along megakaryocyte differentiation. **B)** Transcription factor regulon activity along megakaryocyte differentiation pseudotime. **C)** Gene expression changes along megakaryocyte differentiation pseudotime. **D-F)** Characterization of Eo/Baso/Mast differentiation. **D)** Pseudotime score along Eo/Baso/Mast differentiation. **E)** Transcription factor regulon activity along Eo/Baso/Mast differentiation pseudotime. **F)** Gene expression changes along Eo/Baso/Mast differentiation pseudotime. **G-I)** Characterization of neutrophil differentiation. **G)** Pseudotime score along neutrophil differentiation. **H)** Transcription factor regulon activity along neutrophil differentiation pseudotime **I)** Gene expression changes along neutrophil differentiation pseudotime

**Supplemental Fig 7. Characterization of broad differentiation stages implicated in AML**
**A)** Correlation of the CLR-normalized relative abundance of each projected cell state along the hematopoietic hierarchy, spanning 318 patient samples. **B**) Consolidation of precise cell states into broad differentiation stages. NMF was used to identify groups of correlated cell states, and the top-weighted cell states driving each NMF component were combined into broad differentiation stages. Weights for each cell state underlying each NMF-defined differentiation stage are shown in the heatmap. **C-D**) Enrichment of normal hematopoietic **C**) cell state programs and **D**) cell activity programs within each AML differentiation stage, depicted through AUC values.

**Supplemental Fig 8. Expression of therapeutic targets and LSC signatures by AML differentiation stage**
**A**) Differential expression (DE) results from one-vs-all comparisons between differentiation stages. DE was performed using pseudo-bulk profiles for each differentiation stage from each of the 318 patient samples. Results for therapeutic targets in AML are shown. The log fold change (logFC) statistic for each gene is depicted through the size and color intensity of each dot,

wherein red denotes higher expression while blue denotes lower expression. For visualization, logFC values were capped between -2 and 2. DE results with P < 0.05 are shown, those with FDR < 0.05 are starred. **B-D)** Leukemia stem cell (LSC) signatures across AML differentiation stages. Cells belonging to each differentiation stage from each patient were pooled into pseudo-bulk profiles and LSC signatures were scored by GSVA. Scores are shown for **B)** genes enriched in functional LSC fractions from Ng *et al* 2016 and **C)** Quiescent LSPC signature from Zeng *et al* 2022. **D)** Projection of prior AML cell type annotations from van Galen *et al* 2019 and Zeng *et al* 2022. Labeled cells from three datasets (van Galen *et al* 2019, Abbas *et al* 2021, Wang *et al* 2024) belonging to each AML cell type (LSPC-Quiescent, LSPC-Primed, LSPC-Cycle, GMP-like, ProMono-like, Mono-like, and cDC-like) were collapsed into a centroid for each patient sample. The number of cells that comprise each centroid are depicted by the point size.

**Supplemental Fig 9. Representative samples from each AML differentiation pattern**
For each AML subtype defined from cell state composition analysis, projection results for each of three additional representative samples are depicted. Patient ID and originating dataset are labeled accordingly.

**Supplemental Fig 10. Inference of AML differentiation stage abundance from bulk RNA-seq profiles**
**A**) Depiction of broad differentiation stages in AML, each spanning multiple cell states. **B)** Marker genes specific to each differentiation stage in AML. Cells belonging to each differentiation stage from each patient were pooled prior to differential expression analysis and heatmap visualization. **C)** LASSO regression to estimate AML differentiation stage abundance in 160 pseudo-bulk samples wherein cells from individual patients were aggregated together. Marker genes for each differentiation stage were used as features for LASSO regression, and cross-validation results of inferred abundances by LASSO in held-out patient subsets are depicted for each differentiation stage. **D)** Number of positively and negatively weighted genes comprising each LASSO regression model for predicting abundance of each AML differentiation stage in bulk RNA-seq data. **E)** Predicted and observed differentiation stage abundance across 160 AML pseudo-bulk samples.

**Supplemental Fig 11. Clinical and genomic correlates of AML differentiation stage abundance**
Associations between inferred abundance of AML differentiation stages with genomic alterations across AML patient samples profiled by RNA-seq. The strength of each association, quantified as the test statistic from a generalized linear model adjusting for cohort as a covariate, is depicted through the size and color intensity of each dot, wherein green denotes higher abundance and purple denotes lower abundance. Only associations with unadjusted P < 0.05 are shown, associations with FDR < 0.05 are starred. Associations are depicted for each of **A)** FAB morphology (n=411 patients across TCGA, BeatAML, and Leucegene), **B)** WHO 2022 classification (n=1166 across 5 cohorts re-analyzed by Severens *et al* 2024, consisting of TCGA, BeatAML2, Leucegene, TARGET, and LUMC), **C)** ICC 2022 classification (n=1192 patients across 5 cohorts re-analyzed by Severens *et al* 2024), **D)** Hierarchy classification from Zeng *et al* 2022 (n=864 patients spanning TCGA, BeatAML, and Leucegene), **E)** Gene

expression clusters from Severens *et al* 2024 (n=1224 patients across 5 cohorts re-analyzed by Severens *et al* 2024), **F)** Integrated RNA + Hierarchy + Methylation based classification from Ma *et al* 2024 (n=812 patients spanning TCGA, BeatAML, and Leucegene), and **G)** Disease history (n=487 patients from BeatAML2). **H-J)** Associations for specific mutation patterns spanning 1224 patients across 5 cohorts re-analyzed by Severens *et al* 2024 (TCGA, BeatAML2, Leucegene, TARGET, LUMC). Comparisons are depicted for **H)** CEBPA mutation type (n=99) versus CEBPA wildtype, **I)** FLT3 mutation type (n=340) versus FLT3 wildtype, **J)** NPM1 mutation combination (n=249 with normal karyotype) versus NPM1 wildtype.

**Supplemental Fig 12. Analysis of engraftment patterns among CD34 and CD38 sorted fractions in AML**

AML cells from 74 primary patient samples from Ng et al 2016 were sorted into four fractions based on CD34 and CD38 expression and each fraction was transplanted into NSG mice at varying cell doses. Injected right femur engraftment at 12-weeks post-transplant was evaluated by flow cytometry. Fractions that initiated CD45+CD33+ leukemic grafts were scored as LSC+ (dark blue), those that did not initiate grafts were scored as LSC- (light grey), and those that initiated grafts with both CD33+ myeloid and CD19+ lymphoid cells were scored as multilineage (green) and presumed to be normal or pre-leukemic. Patient samples were grouped together based on engraftment patterns across their CD34 and CD38 fractions as well as AML hierarchy composition by deconvolution of bulk RNA-seq profiles (Zeng et al 2022) to identify subsets with potential involvement of multiple LSC populations.

**Supplemental Fig 13. Cell state classification in AML 90240 primary and xenograft samples**

**A)** UMAP embedding and AML cell type assignments of scRNA-seq from primary samples of AML 90240. **B)** UMAP embedding and AML cell type assignments from patient-derived xenograft (PDX) samples of AML 90240. **C)** Density plot of unsorted bulk, CD34+CD38- sorted, and CD34-CD38+ sorted primary fractions from AML 90240. **D)** Density plot of PDX generated from CD34+CD38- primary cells and PDX generated from CD34-CD38+ primary cells from AML 90240. **E)** BoneMarrowMap projection of primary AML fractions from (C). **F)** BoneMarrowMap projection of xenografted AML cells from (D).

**Supplemental Fig 14. Distinct LSC-driven leukemia cell hierarchies co-existing within AML 90394**

**A-F)** Experimental workflow for identifying co-existing LSC-driven hierarchies as in Figure 6, applied to AML #90394. **A)** Immunophenotype and scRNA-seq cell state composition of primary AML. **B)** scRNA-seq projection results of primary AML sample. **C)** Immunophenotype and scRNA-seq composition for CD34+CD38- fraction, representing 0.17% of primary cells. **D)** Immunophenotype and scRNA-seq composition for PDX derived from CD34+CD38- fraction, recapitulating the primary AML hierarchy. **E)** Immunophenotype and scRNA-seq composition for CD34-CD38+ fraction, representing 16% of primary cells. **F)** Immunophenotype and scRNA-seq composition for PDX derived from CD34-CD38+ fraction, restricted to mature AML populations.

**Supplemental Fig 15. Cell state classification in AML 90394 primary and xenograft samples**

**A)** UMAP embedding and AML cell type assignments of scRNA-seq from primary samples of AML 90394. **B)** UMAP embedding and AML cell type assignments from patient-derived xenograft (PDX) samples of AML 90394. **C)** Density plot of unsorted bulk, CD34+CD38- sorted, and CD34-CD38+ sorted primary fractions from AML 90394. **D)** Density plot of PDX generated from CD34+CD38- primary cells and PDX generated from CD34-CD38+ primary cells from AML 90394. **E)** BoneMarrowMap projection of primary AML fractions from (C). **F)** BoneMarrowMap projection of xenografted AML cells from (D).

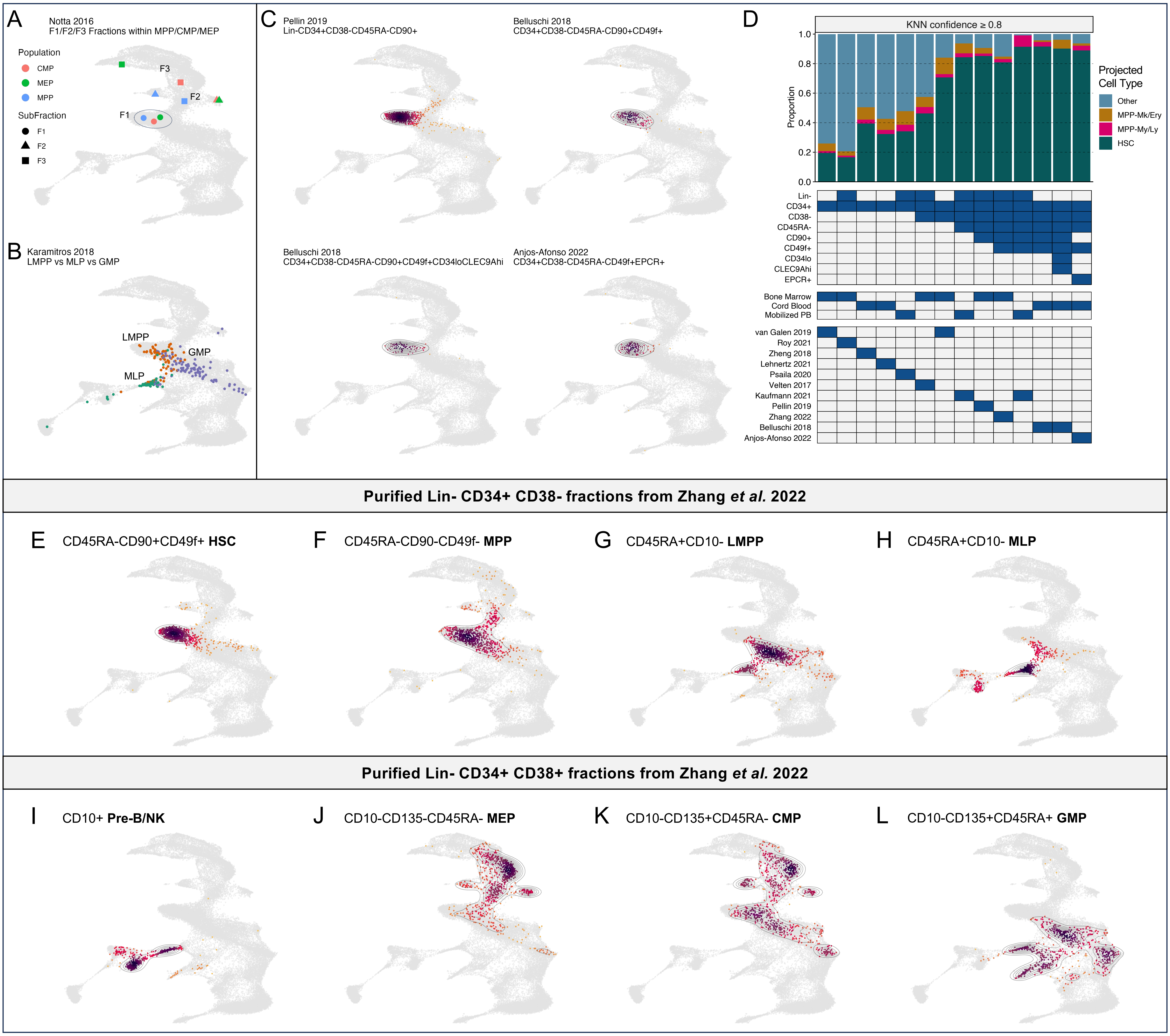

**Supplemental Figure 1. Projection of sorted HSPC populations onto BoneMarrowMap**

**A)** Projection of purified multilineage (F1) and Mk/Ery restricted (F2, F3) sub-fractions of MPP, CMP, and MEP from Notta *et al* 2016. **B)** Projection of scRNA-seq profiles purified with functionally informed gating of LMPP, GMP, and MLP populations from Karamitros *et al* 2018. **C)** Projection of scRNA-seq profiles from HSCs purified using different markers of human HSCs. **D)** Proportion of transcriptional HSCs and MPPs projected from scRNA-seq data sorted with varying levels of immunophenotypic purity. The combination of markers used, tissue source, and dataset name are also depicted. **E-L)** Projection of scRNA-seq profiles of immunophenotypically defined HSPC populations from Zhang *et al* 2022, spanning **E)** HSC, **F)** MPP, **G)** LMPP, **H)** MLPs, **I)** Pre-B/NK, **J)** MEP, **K)** CMP, and **L)** GMP.

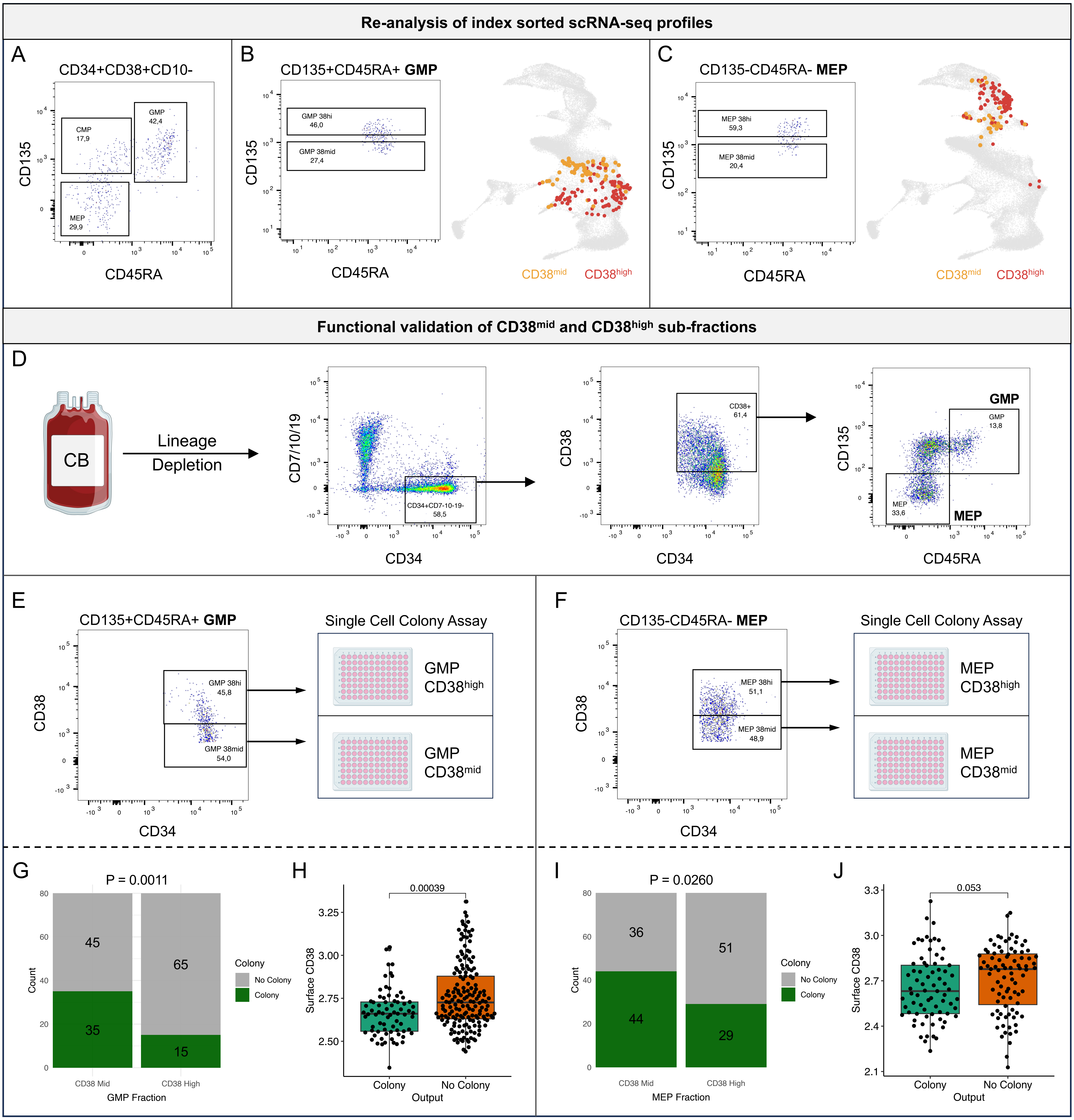

**Supplemental Figure 2. Functionally distinct CD38 fractions within human GMPs and MEPs**

**A-C)** Re-analysis of index-sorted scRNA-seq profiles from Velten *et al* 2017. **A)** Gating of MEP, CMP, and GMPs within CD34+CD38+CD10- progenitors. **B)** Gating and scRNA-seq projection of CD38-mid and CD38-high GMP fractions. **C)** Gating and scRNA-seq projection of CD38-mid and CD38-high MEP fractions. **D-F)** Single-cell colony assays with index-sorting of CD38-mid and CD38-high **E)** GMP fractions and **F)** MEP fractions from umbilical cord blood. **G)** Single-cell colony formation outcomes within CD38-mid and CD38-high GMPs and **H)** surface CD38 levels of individual GMPs that did and did not form colonies. **I)** Single-cell colony formation outcomes within CD38-mid and CD38-high MEPs and **J)** surface CD38 levels of individual MEPs that did and did not form colonies.

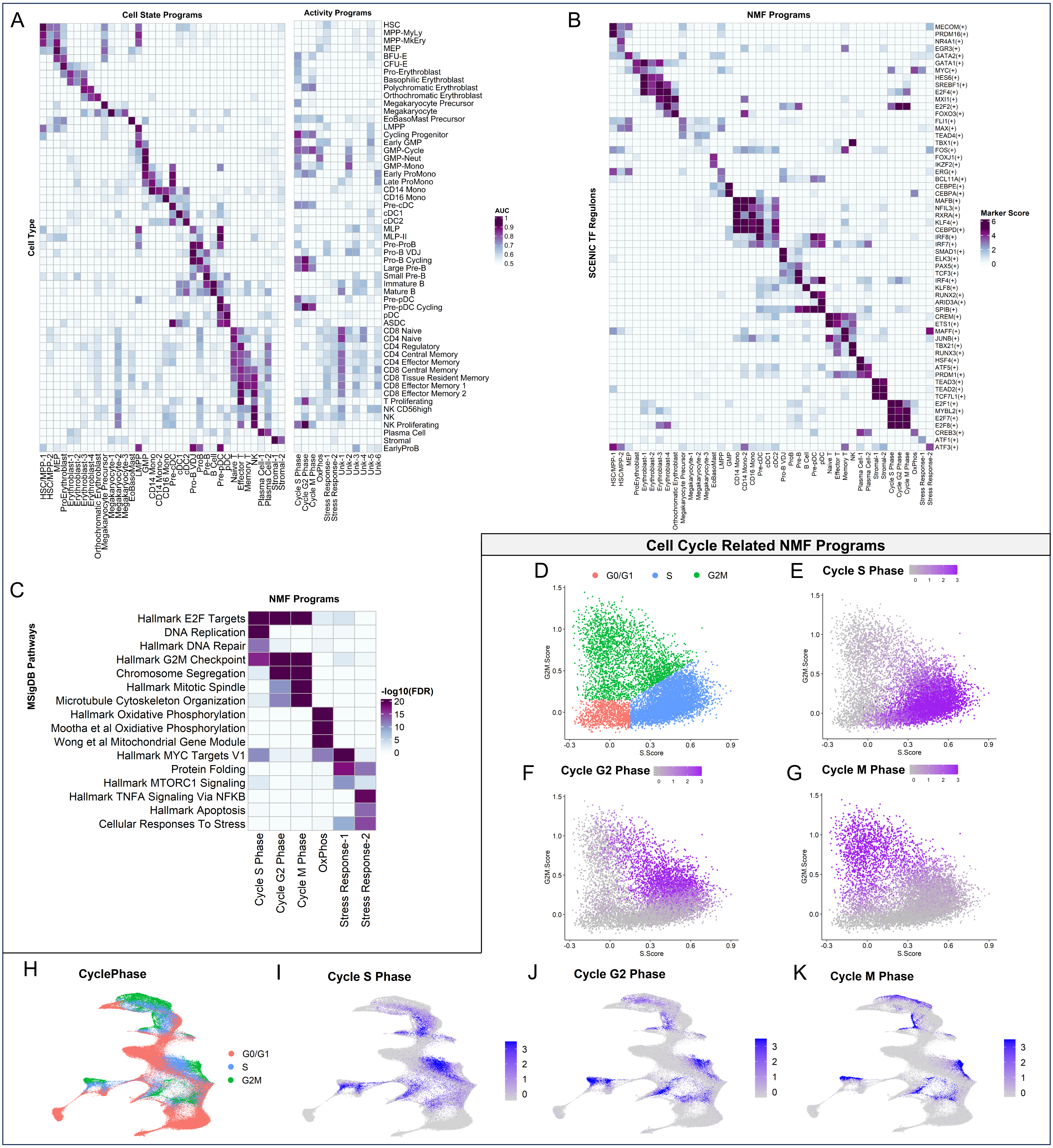

**Supplemental Figure 3. Identification of gene expression programs through consensus NMF**
**A)** Unsupervised discovery of 48 gene expression programs (GEP) across human hematopoiesis by cNMF. 36 GEPs correspond to hematopoietic cell states (cell state programs) and 12 GEPs correspond to other cellular processes (activity programs). The relative activity of each program within each hematopoietic cell state is depicted through a heatmap of AUC values. **B)** Transcription factor (TF) regulons associated with each GEP based on a composite score representing both the correlation of GEPs with each TF regulon score as well as the significance of overlap between genes defining each GEP and each TF regulon. **C)** Key pathways significantly associated with activity GEPs. **D-G)** Cell cycle plot of S vs G2M scores within HSPCs, depicting **D)** cell cycle assignments and **E)** S-phase, **F)** G2-phase, and **G)** M-phase program expression. **H-I)** UMAP of hematopoietic cell states depicting **H)** cell cycle assignments and **I)** S-phase, **J)** G2-phase, and **K)** M-phase program expression.

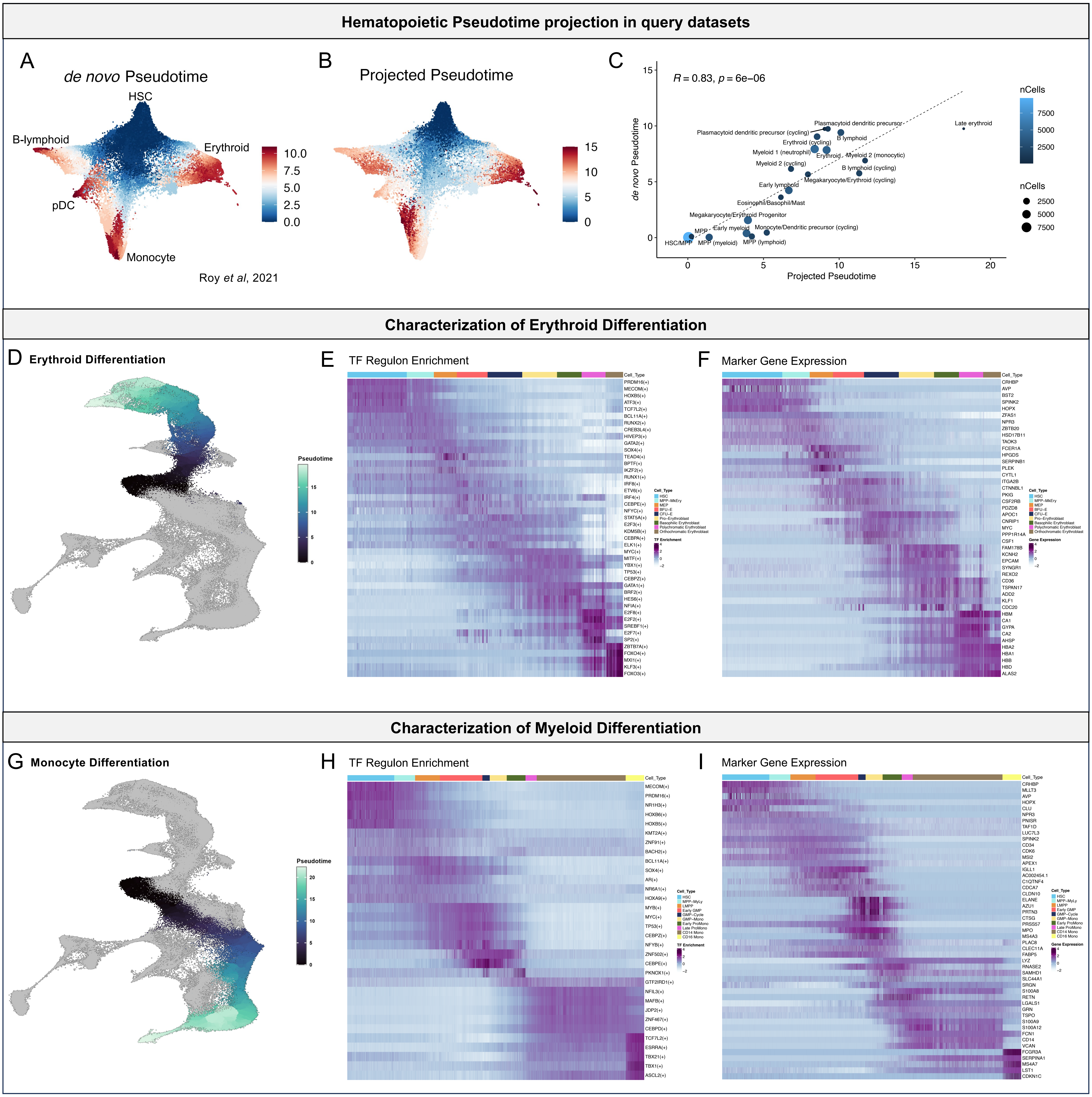

**Supplemental Figure 4. Pseudotime analysis and characterization of erythroid and monocytic differentiation A-C)** Projection and validation of hematopoietic pseudotime values in query data from Roy *et al* 2021. **A)** Pseudotime values calculated *de novo* on HSPC transcriptomes from Roy *et al* 2021 using Monocle3. **B)** Pseudotime values inferred through BoneMarrowMap projection of HSPC transcriptomes from Roy *et al* 2021. **C)** Comparison of *de novo* and projected pseudotime values across cell states annotated in Roy *et al* 2021. Average values for each cell state are depicted and the size and color of each cell state correspond to the number of cells within that state. **D-F)** Characterization of erythroid differentiation. **D)** Pseudotime score along erythroid differentiation. **E)** Transcription factor regulon activity along erythroid differentiation pseudotime. **F)** Gene expression changes along erythroid differentiation pseudotime. **G-I)** Characterization of myeloid differentiation. **G)** Pseudotime score along monocyte differentiation. **H)** Transcription factor regulon activity along monocyte differentiation pseudotime. **I)** Gene expression changes along monocyte differentiation pseudotime.

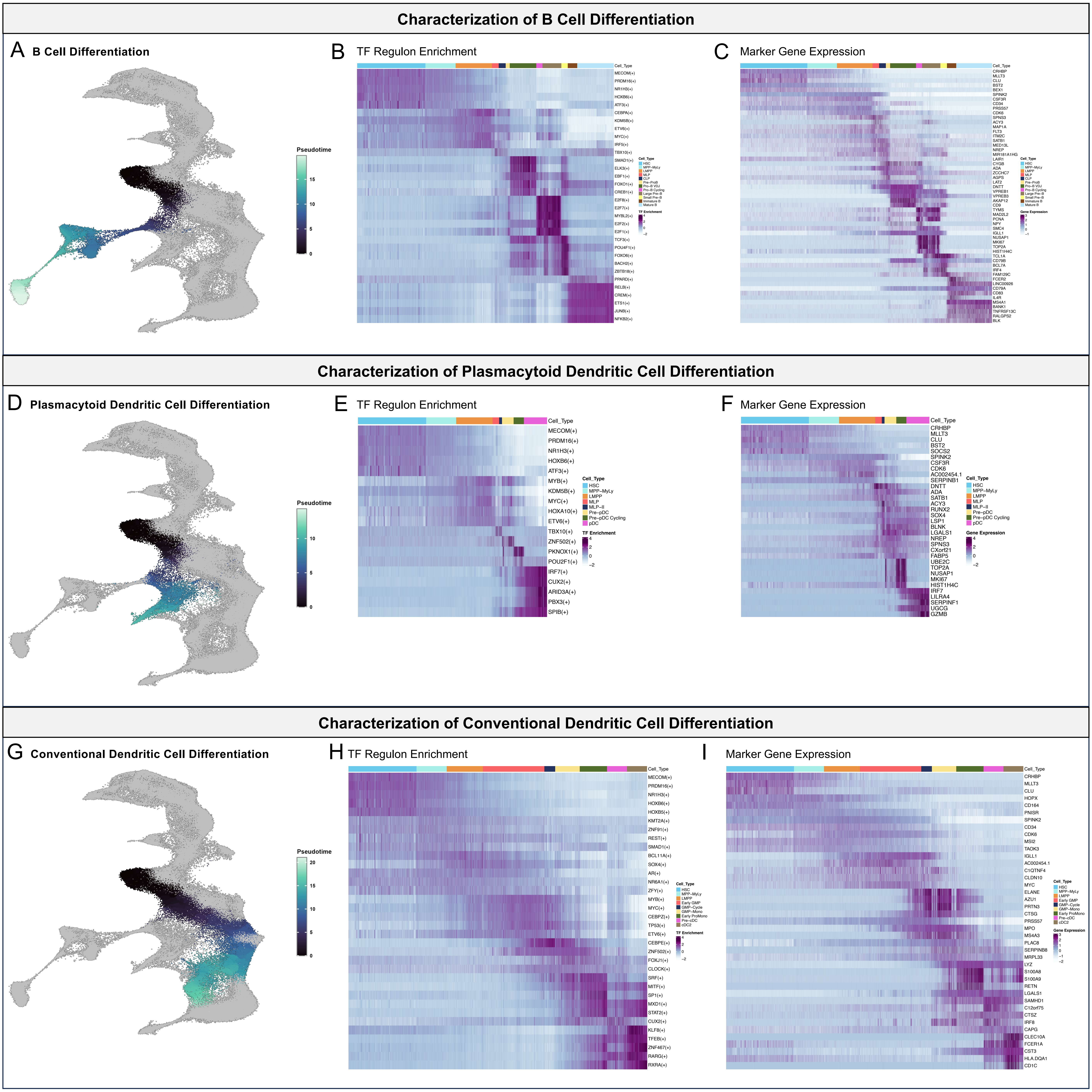

**Supplemental Figure 5. Characterization of B-lymphoid and dendritic cell differentiation**
**A-C)** Characterization of B cell differentiation. **A)** Pseudotime score along B cell differentiation. **B)** Transcription factor regulon activity along B cell differentiation pseudotime. **C)** Gene expression changes along B cell differentiation pseudotime. **D-F)** Characterization of plasmacytoid dendritic cell (pDC) differentiation. **D)** Pseudotime score along pDC differentiation. **E)** Transcription factor regulon activity along pDC differentiation pseudotime. **F)** Gene expression changes along pDC differentiation pseudotime. **G-I)** Characterization of conventional dendritic cell (cDC) differentiation. **G)** Pseudotime score along cDC differentiation. **H)** Transcription factor regulon activity along cDC differentiation pseudotime. **I)** Gene expression changes along cDC differentiation pseudotime.

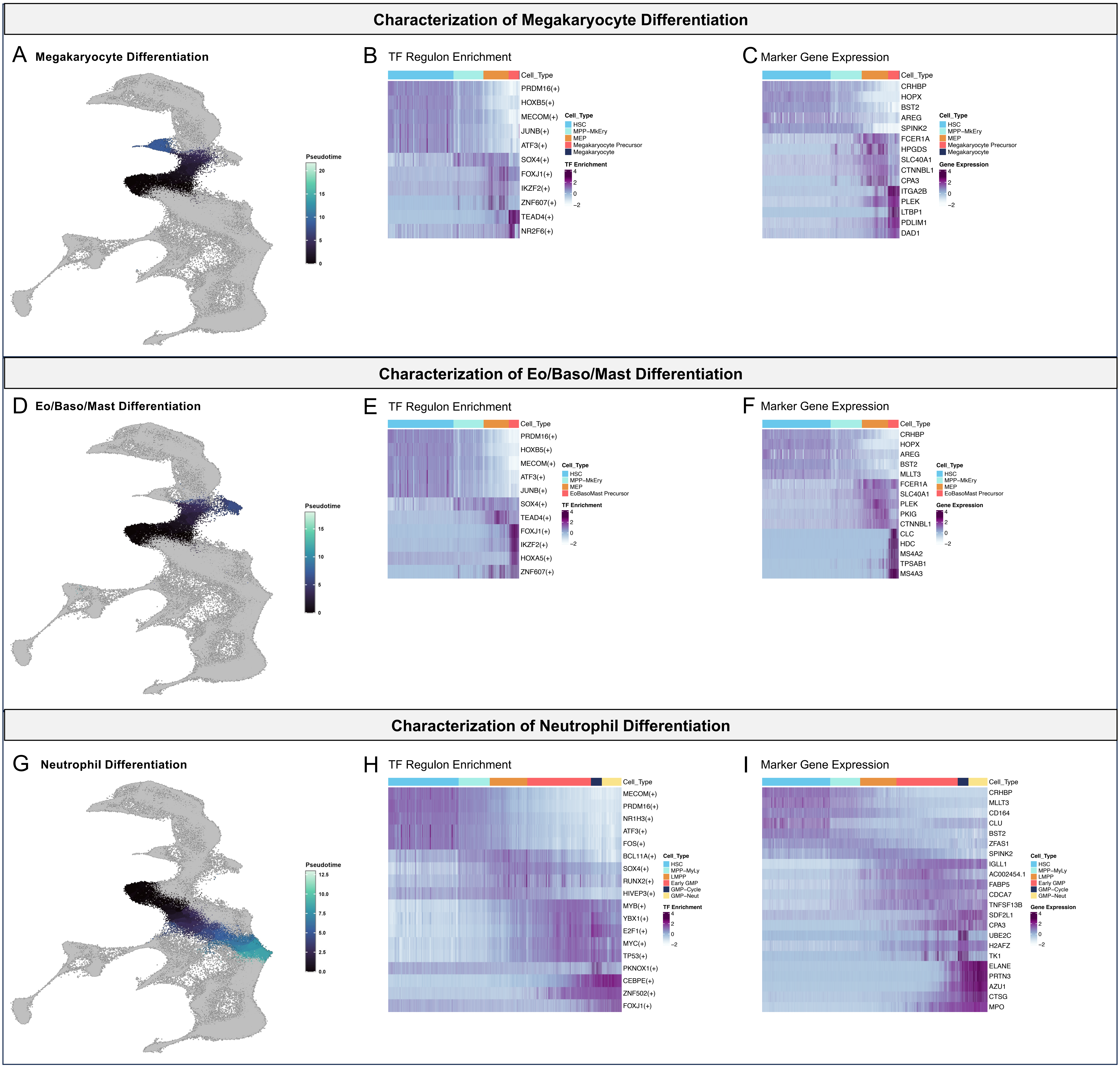

**Supplemental Figure 6. Characterization of megakaryocyte and granulocyte differentiation**
**A-C)** Characterization of megakaryocyte differentiation. **A)** Pseudotime score along megakaryocyte differentiation. **B)** Transcription factor regulon activity along megakaryocyte differentiation pseudotime. **C)** Gene expression changes along megakaryocyte differentiation pseudotime. **D-F)** Characterization of Eo/Baso/Mast differentiation. **D)** Pseudotime score along Eo/Baso/Mast differentiation. **E)** Transcription factor regulon activity along Eo/Baso/Mast differentiation pseudotime. **F)** Gene expression changes along Eo/Baso/Mast differentiation pseudotime. **G-I)** Characterization of neutrophil differentiation. **G)** Pseudotime score along neutrophil differentiation. **H)** Transcription factor regulon activity along neutrophil differentiation pseudotime. **I)** Gene expression changes along neutrophil differentiation pseudotime

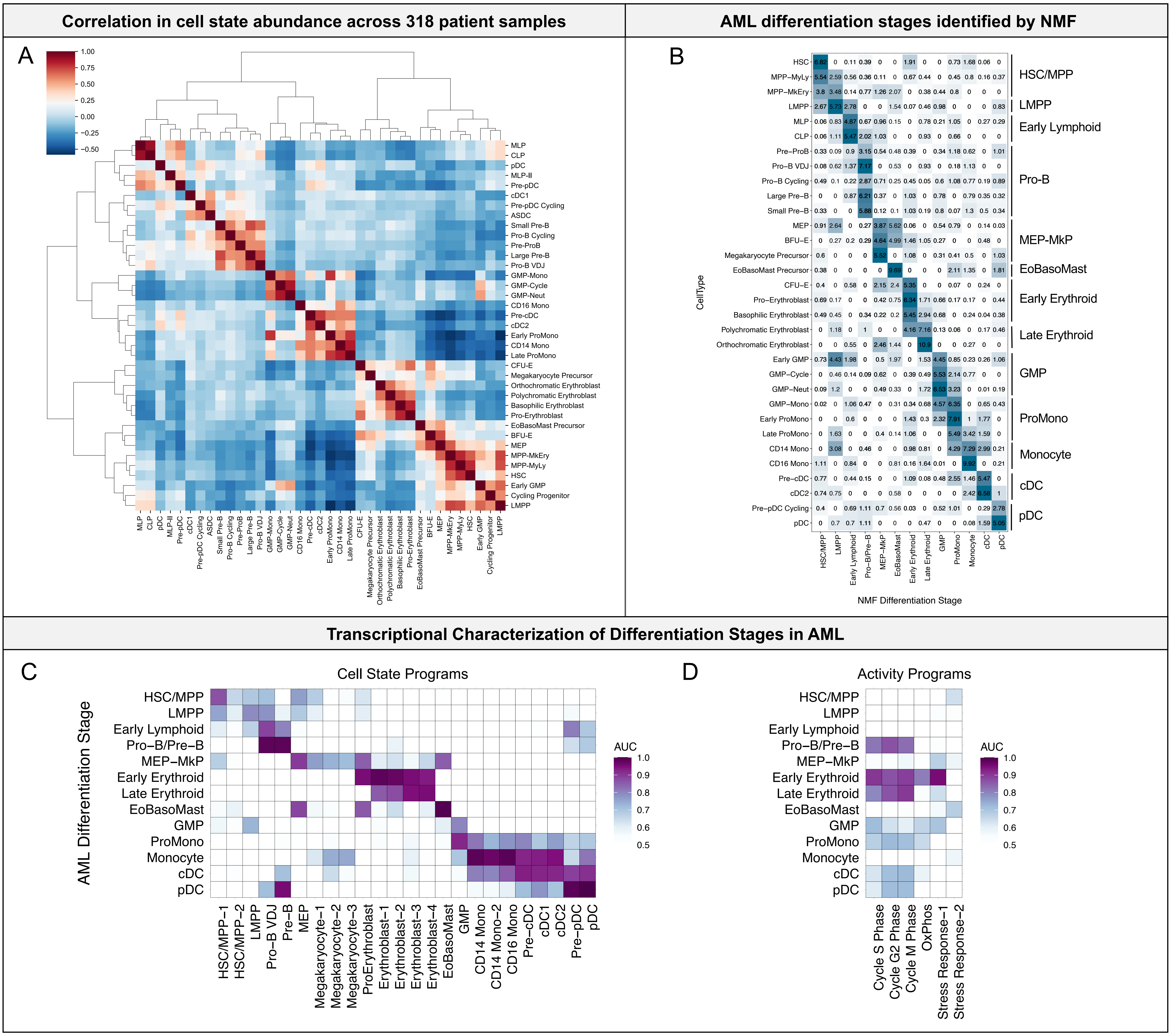

**Supplemental Figure 7. Characterization of broad differentiation stages implicated in AML**

**A)** Correlation of the CLR-normalized relative abundance of each projected cell state along the hematopoietic hierarchy, spanning 318 patient samples. **B**) Consolidation of precise cell states into broad differentiation stages. NMF was used to identify groups of correlated cell states, and the top-weighted cell states driving each NMF component were combined into broad differentiation stages. Weights for each cell state underlying each NMF-defined differentiation stage are shown in the heatmap. **C-D**) Enrichment of normal hematopoietic **C**) cell state programs and **D**) cell activity programs within each AML differentiation stage, depicted through AUC values.

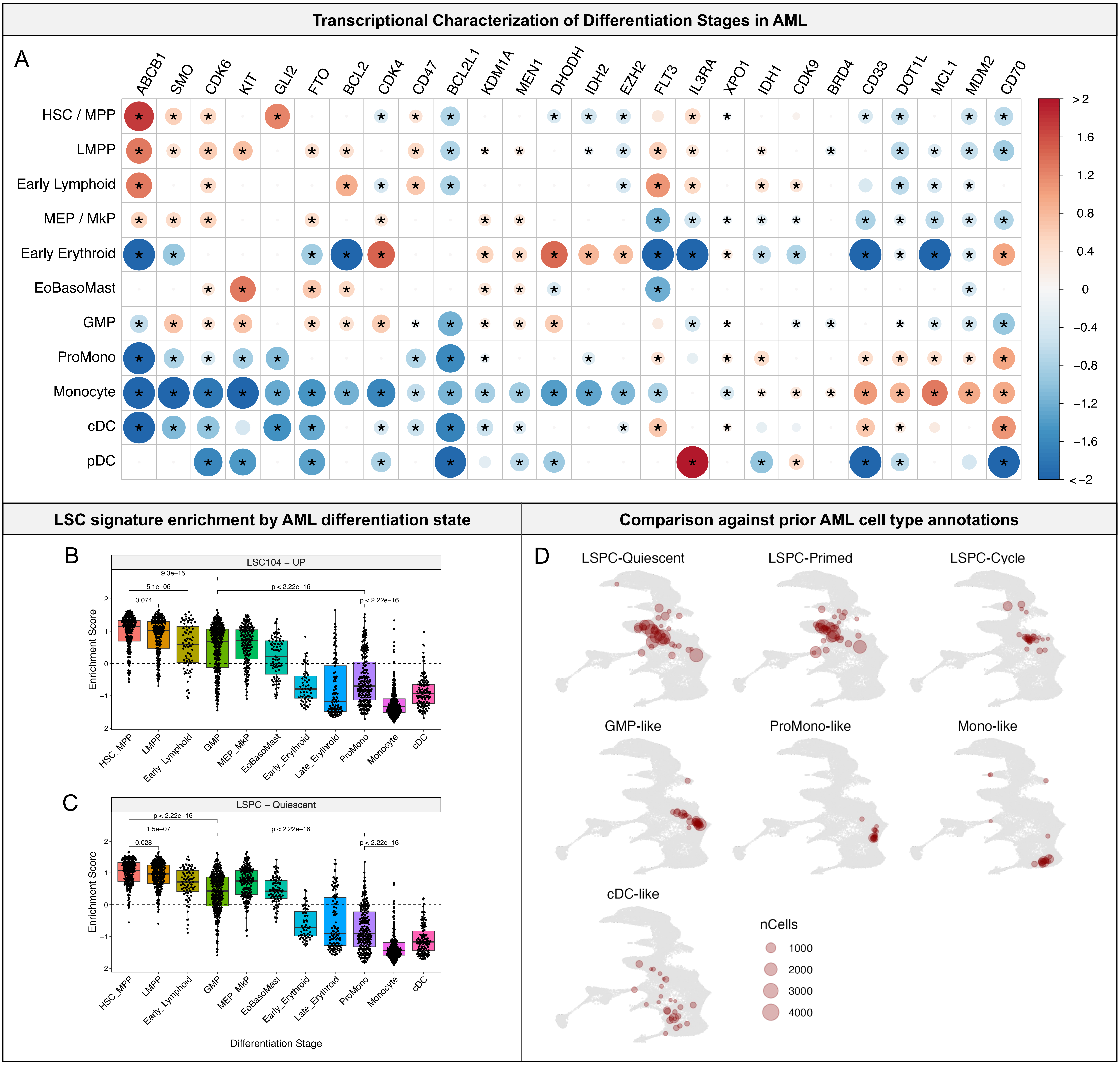

**Supplemental Figure 8. Expression of therapeutic targets and LSC signatures by AML differentiation stage**
**A**) Differential expression (DE) results from one-vs-all comparisons between differentiation stages. DE was performed using pseudo-bulk profiles for each differentiation stage from each of the 318 patient samples. Results for therapeutic targets in AML are shown. The log fold change (logFC) statistic for each gene is depicted through the size and color intensity of each dot, wherein red denotes higher expression while blue denotes lower expression. For visualization, logFC values were capped between -2 and 2. DE results with P < 0.05 are shown, those with FDR < 0.05 are starred. **B-D)** Leukemia stem cell (LSC) signatures across AML differentiation stages. Cells belonging to each differentiation stage from each patient were pooled into pseudo-bulk profiles and LSC signatures were scored by GSVA. Scores are shown for **B)** genes enriched in functional LSC fractions from Ng *et al* 2016 and **C)** Quiescent LSPC signature from Zeng *et al* 2022. **D)** Projection of prior AML cell type annotations from van Galen *et al* 2019 and Zeng *et al* 2022. Labeled cells from three datasets (van Galen *et al* 2019, Abbas *et al* 2021, Wang *et al* 2024) belonging to each AML cell type (LSPC-Quiescent, LSPC-Primed, LSPC-Cycle, GMP-like, ProMono-like, Mono-like, and cDC-like) were collapsed into a centroid for each patient sample. The number of cells that comprise each centroid are depicted by the point size.

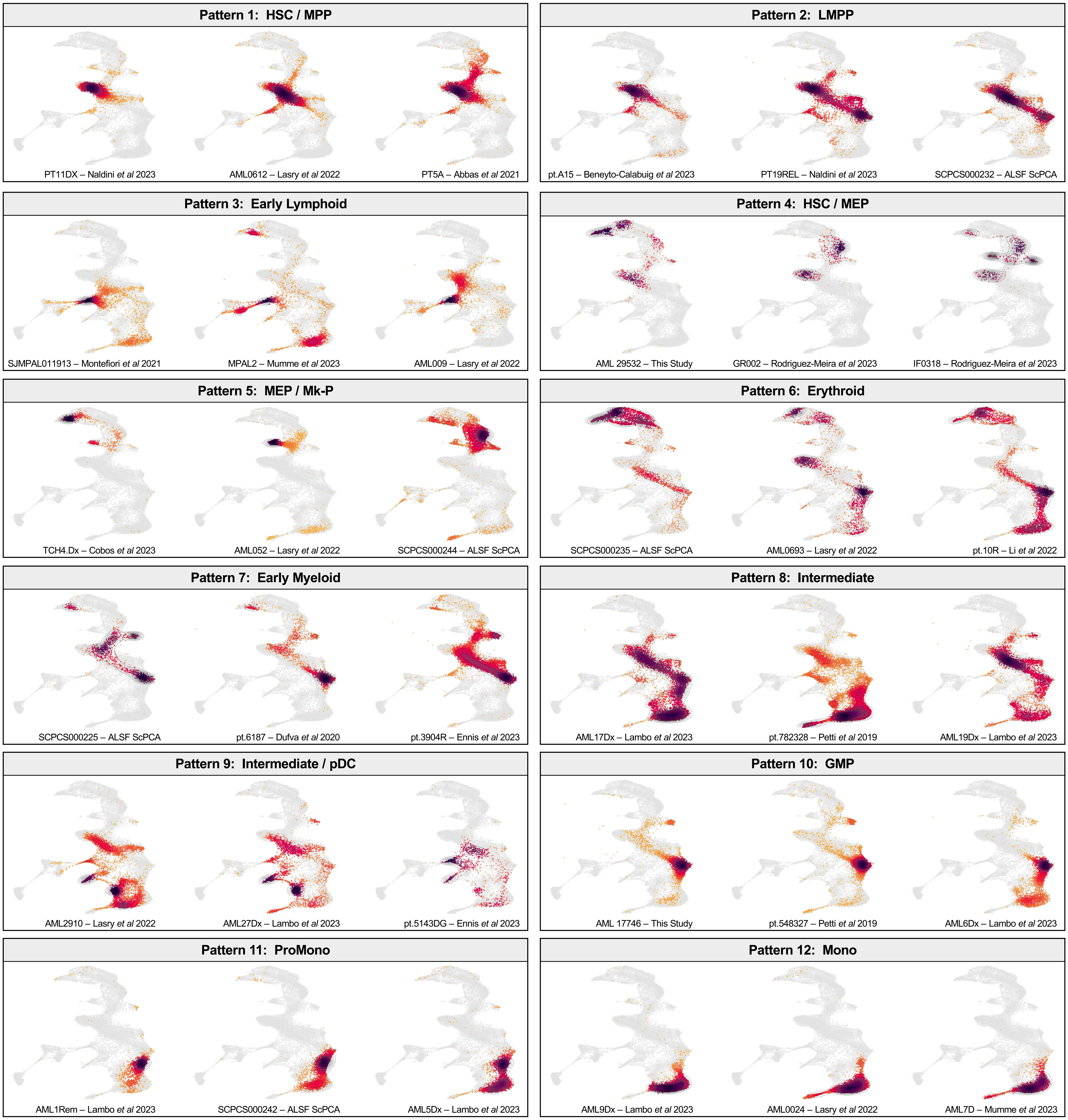

**Supplemental Figure 9. Representative samples from each AML differentiation pattern**
For each AML differentiation pattern defined from cell state composition analysis, projection results for each of three additional representative scRNA-seq samples are depicted. Patient ID and originating dataset are labeled accordingly.

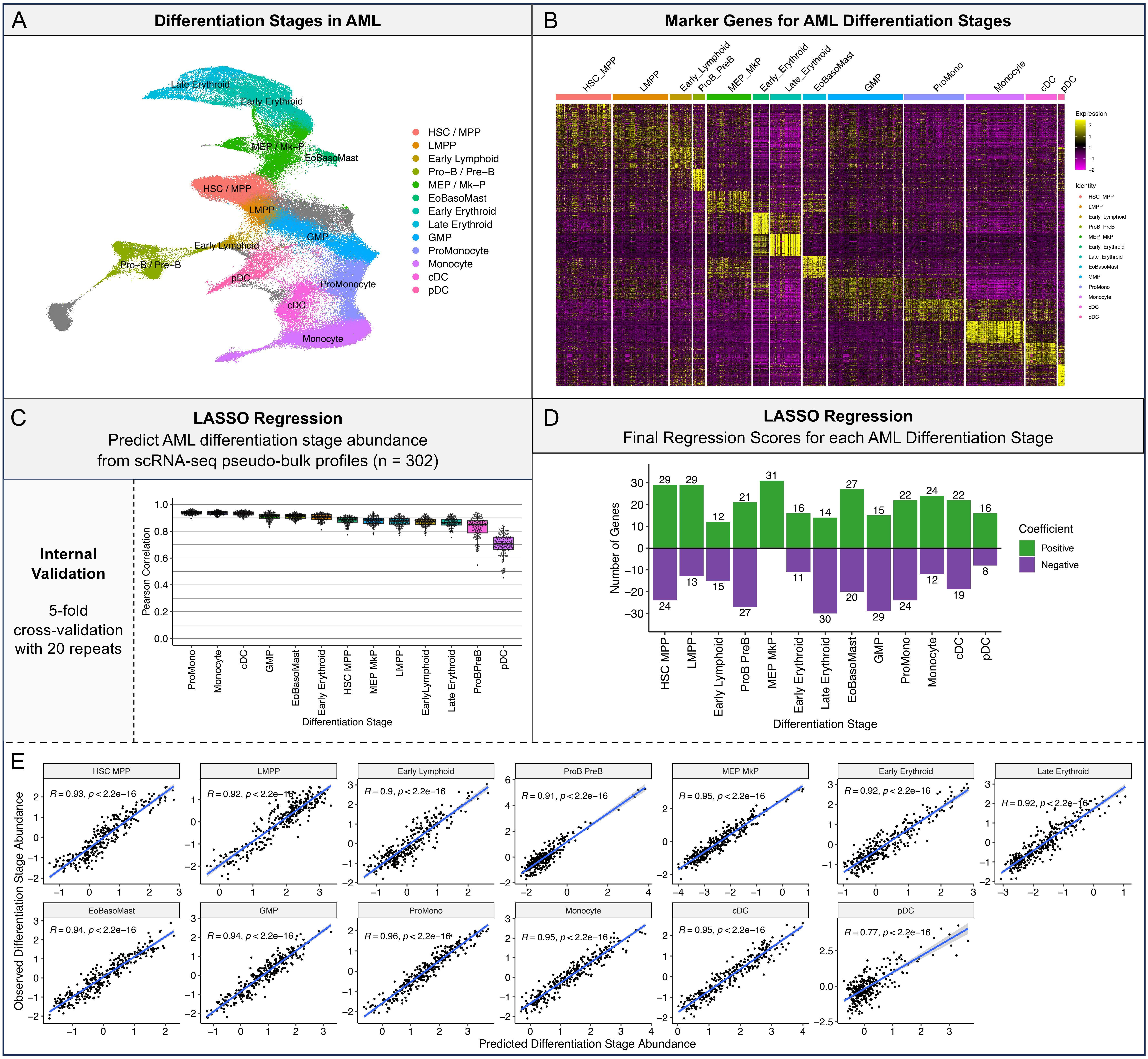

**Supplemental Figure 10. Inference of AML differentiation stage abundance from bulk RNA-seq profiles**
**A**) Depiction of broad differentiation stages in AML, each spanning multiple cell states. **B)** Marker genes specific to each differentiation stage in AML. Cells belonging to each differentiation stage from each patient were pooled prior to differential expression analysis and heatmap visualization. **C)** LASSO regression to estimate AML differentiation stage abundance in 302 pseudo-bulk samples wherein cells from individual patients were aggregated together. Marker genes for each differentiation stage were used as features for LASSO regression, and cross-validation results of inferred abundances by LASSO in held-out patient subsets are depicted for each differentiation stage. **D)** Number of positively and negatively weighted genes comprising each LASSO regression model for predicting abundance of each AML differentiation stage in bulk RNA-seq data. **E)** Predicted and observed differentiation stage abundance across 302 AML pseudo-bulk samples.

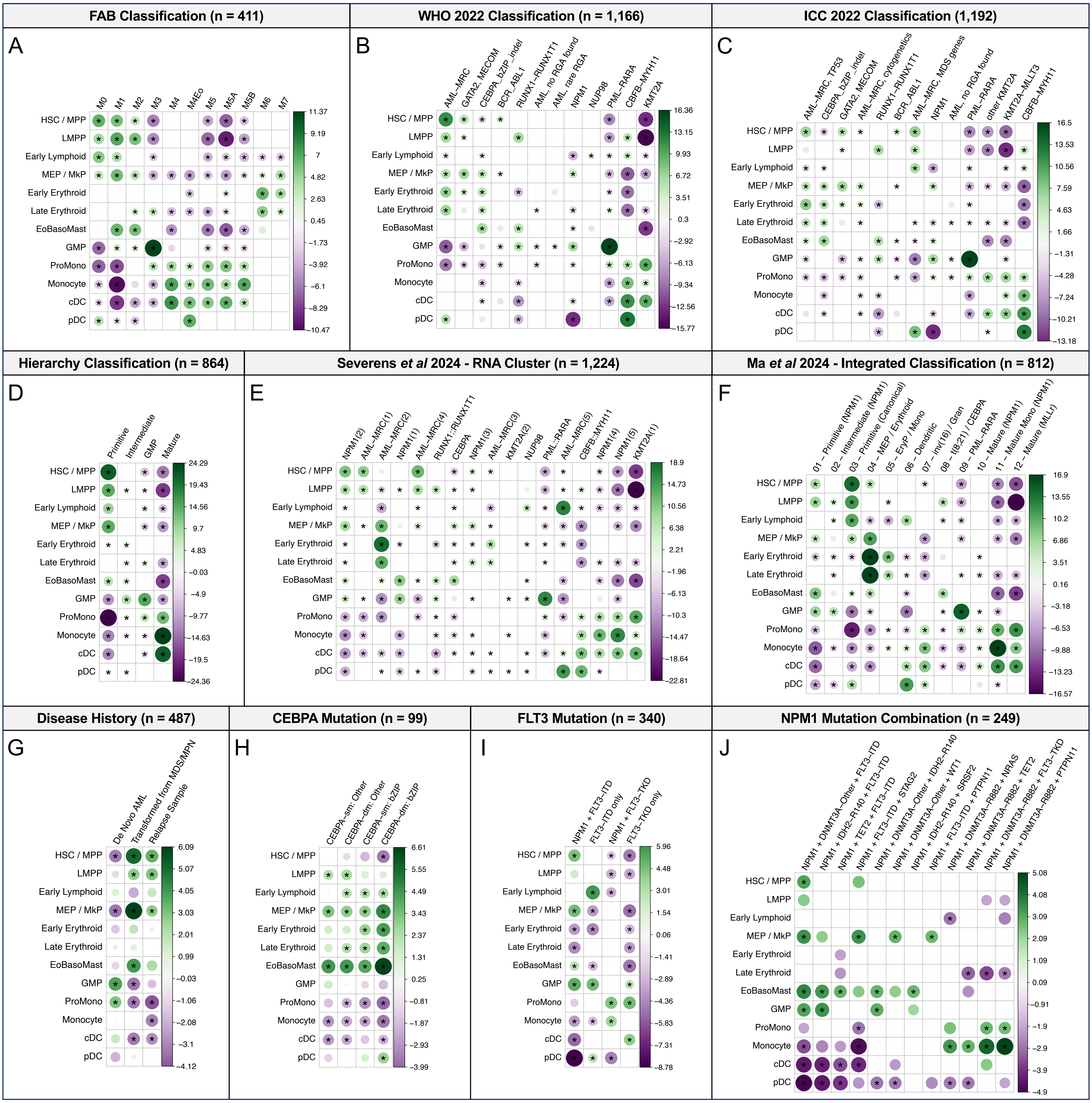

**Supplemental Figure 11. Clinical and genomic correlates of AML differentiation stage abundance**

Associations between inferred abundance of AML differentiation stages with genomic alterations across AML patient samples profiled by RNA-seq. The strength of each association, quantified as the test statistic from a generalized linear model adjusting for cohort as a covariate, is depicted through the size and color intensity of each dot, wherein green denotes higher abundance and purple denotes lower abundance. Only associations with unadjusted P < 0.05 are shown, associations with FDR < 0.05 are starred. Associations are depicted for each of **A)** FAB morphology (n=411 patients across TCGA, BeatAML, and Leucegene), **B)** WHO 2022 classification (n=1166 across 5 cohorts re-analyzed by Severens *et al* 2024, consisting of TCGA, BeatAML2, Leucegene, TARGET, and LUMC), **C)** ICC 2022 classification (n=1192 patients across 5 cohorts re-analyzed by Severens *et al* 2024), **D)** Hierarchy classification from Zeng *et al* 2022 (n=864 patients spanning TCGA, BeatAML, and Leucegene), **E)** Gene expression clusters from Severens *et al* 2024 (n=1224 patients across 5 cohorts re-analyzed by Severens *et al* 2024), **F)** Integrated RNA + Hierarchy + Methylation based classification from Ma *et al* 2024 (n=812 patients spanning TCGA, BeatAML, and Leucegene), and **G)** Disease history (n=487 patients from BeatAML2). **H-J)** Associations for specific mutation patterns spanning 1224 patients across 5 cohorts re-analyzed by Severens *et al* 2024 (TCGA, BeatAML2, Leucegene, TARGET, LUMC). Comparisons are depicted for **H)** CEBPA mutation type (n=99) versus CEBPA wildtype, **I)** FLT3 mutation type (n=340) versus FLT3 wildtype, **J)** NPM1 mutation combination (n=249 with normal karyotype) versus NPM1 wildtype.

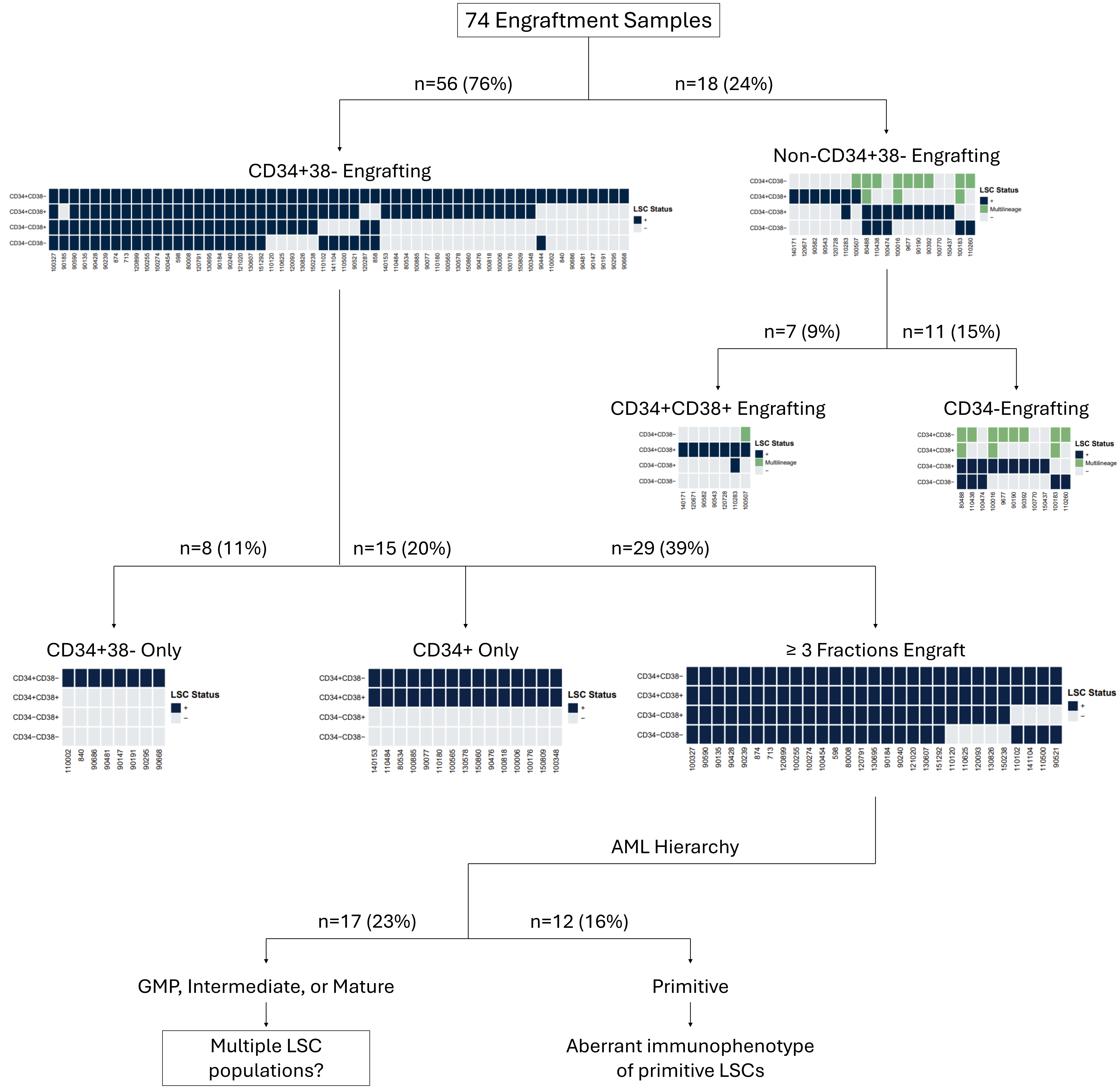

**Supplemental Figure 12. Analysis of engraftment patterns among CD34 and CD38 sorted fractions in AML**
AML cells from 74 primary patient samples from Ng *et al* 2016 were sorted into four fractions based on CD34 and CD38 expression and each fraction was transplanted into NSG mice at varying cell doses. Injected right femur engraftment at 12-weeks post-transplant was evaluated by flow cytometry. Fractions that initiated CD45+CD33+ leukemic grafts were scored as LSC+ (dark blue), those that did not initiate grafts were scored as LSC- (light grey), and those that initiated grafts with both CD33+ myeloid and CD19+ lymphoid cells were scored as multilineage (green) and presumed to be normal or pre-leukemic. Patient samples were grouped together based on engraftment patterns across their CD34 and CD38 fractions as well as AML hierarchy composition by deconvolution of bulk RNA-seq profiles (Zeng *et al* 2022) to identify subsets with potential involvement of multiple LSC populations.

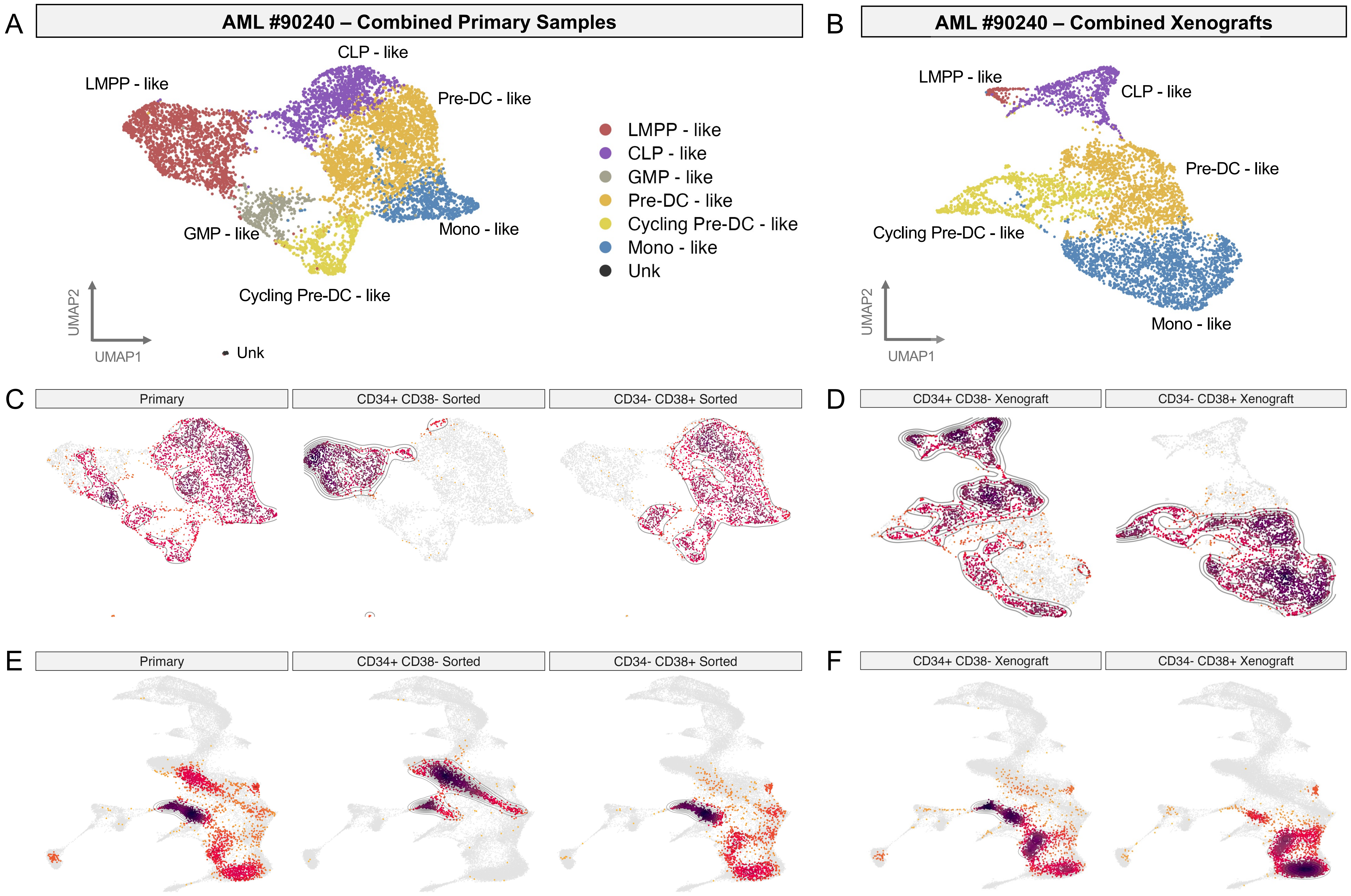

**Supplemental Figure 13. Cell state classification in AML 90240 primary and xenograft samples**
**A)** UMAP embedding and AML cell type assignments of scRNA-seq from primary samples of AML 90240. **B)** UMAP embedding and AML cell type assignments from patient-derived xenograft (PDX) samples of AML 90240. **C)** Density plot of unsorted bulk, CD34+CD38- sorted, and CD34-CD38+ sorted primary fractions from AML 90240. **D)** Density plot of PDX generated from CD34+CD38- primary cells and PDX generated from CD34-CD38+ primary cells from AML 90240. **E)** BoneMarrowMap projection of primary AML fractions from (C). **F)** BoneMarrowMap projection of xenografted AML cells from (D).

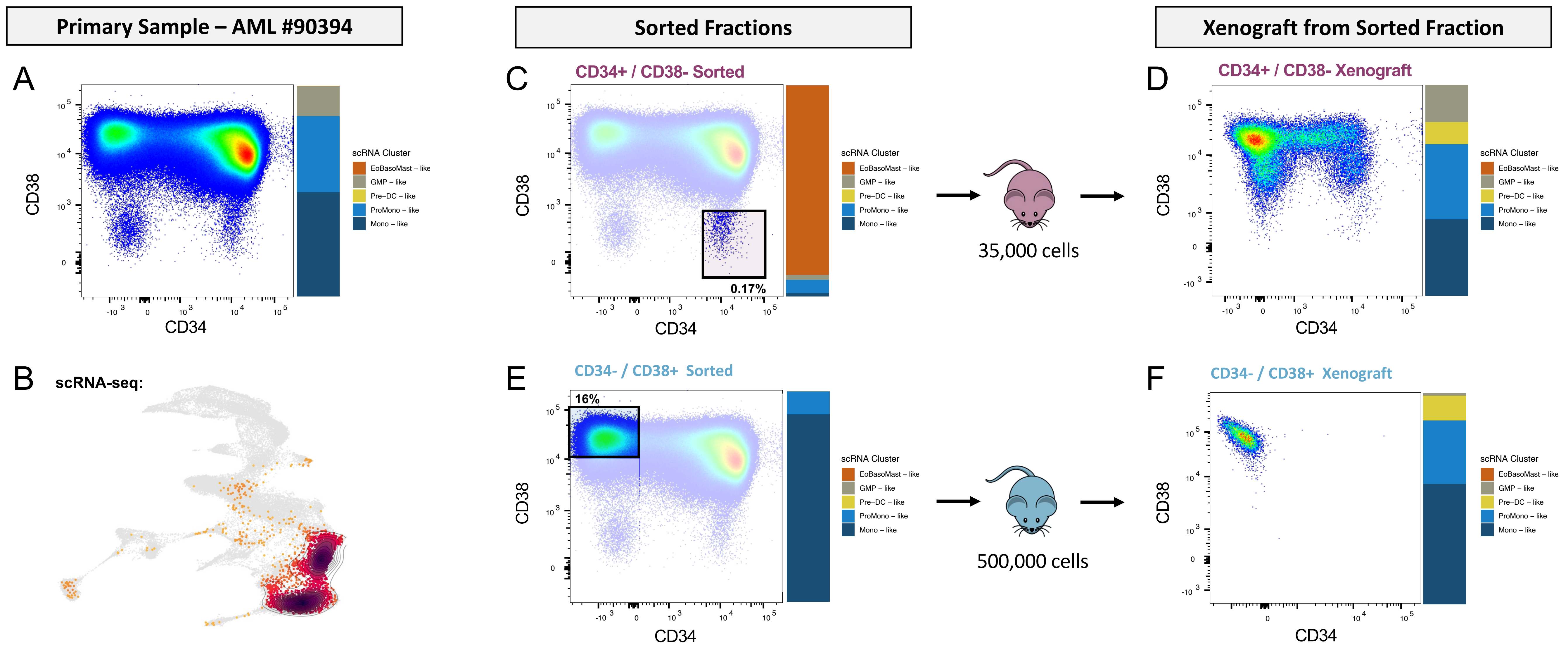

**Supplemental Figure 14. Distinct LSC-driven leukemia cell hierarchies co-existing within AML 90394**
**A-F)** Experimental workflow for identifying co-existing LSC-driven hierarchies as in Figure 6, applied to AML #90394. **A)** Immunophenotype and scRNA-seq cell state composition of primary AML. **B)** scRNA-seq projection results of primary AML sample. **C)** Immunophenotype and scRNA-seq composition for CD34+CD38- fraction, representing 0.17% of primary cells. **D)** Immunophenotype and scRNA-seq composition for PDX derived from CD34+CD38- fraction, recapitulating the primary AML hierarchy. **E)** Immunophenotype and scRNA-seq composition for CD34-CD38+ fraction, representing 16% of primary cells. **F)** Immunophenotype and scRNA-seq composition for PDX derived from CD34-CD38+ fraction, restricted to mature AML populations.

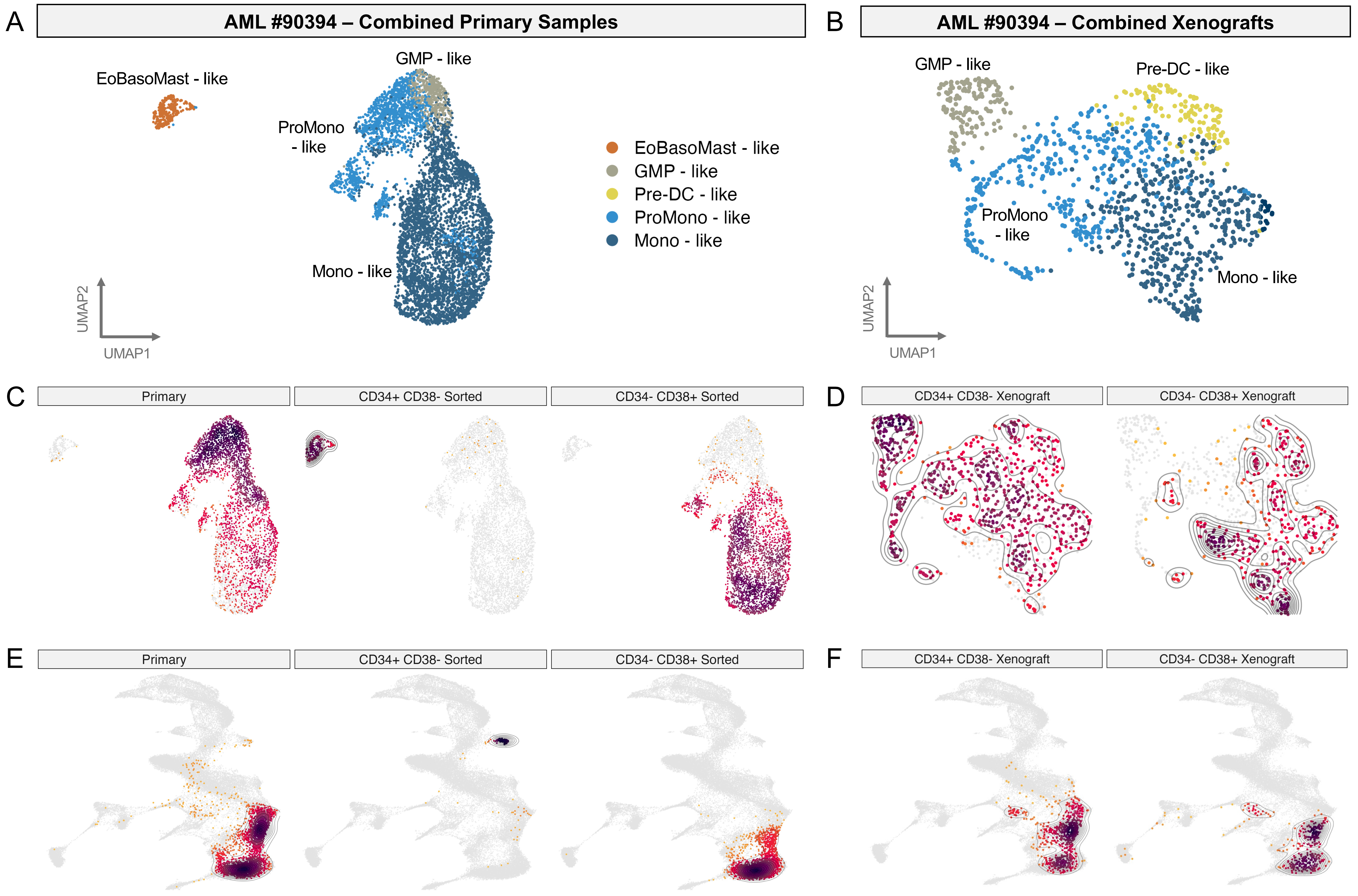

**Supplemental Figure 15. Cell state classification in AML 90394 primary and xenograft samples**
**A)** UMAP embedding and AML cell type assignments of scRNA-seq from primary samples of AML 90394. **B)** UMAP embedding and AML cell type assignments from patient-derived xenograft (PDX) samples of AML 90394. **C)** Density plot of unsorted bulk, CD34+CD38- sorted, and CD34-CD38+ sorted primary fractions from AML 90394. **D)** Density plot of PDX generated from CD34+CD38- primary cells and PDX generated from CD34-CD38+ primary cells from AML 90394. **E)** BoneMarrowMap projection of primary AML fractions from (C). **F)** BoneMarrowMap projection of xenografted AML cells from (D).
